# COVID-19 coronavirus vaccine design using reverse vaccinology and machine learning

**DOI:** 10.1101/2020.03.20.000141

**Authors:** Edison Ong, Mei U Wong, Anthony Huffman, Yongqun He

## Abstract

To ultimately combat the emerging COVID-19 pandemic, it is desired to develop an effective and safe vaccine against this highly contagious disease caused by the SARS-CoV-2 coronavirus. Our literature and clinical trial survey showed that the whole virus, as well as the spike (S) protein, nucleocapsid (N) protein, and membrane (M) protein, have been tested for vaccine development against SARS and MERS. However, these vaccine candidates might lack the induction of complete protection and have safety concerns. We then applied the Vaxign reverse vaccinology tool and the newly developed Vaxign-ML machine learning tool to predict COVID-19 vaccine candidates. By investigating the entire proteome of SARS-CoV-2, six proteins, including the S protein and five non-structural proteins (nsp3, 3CL-pro, and nsp8-10), were predicted to be adhesins, which are crucial to the viral adhering and host invasion. The S, nsp3, and nsp8 proteins were also predicted by Vaxign-ML to induce high protective antigenicity. Besides the commonly used S protein, the nsp3 protein has not been tested in any coronavirus vaccine studies and was selected for further investigation. The nsp3 was found to be more conserved among SARS-CoV-2, SARS-CoV, and MERS-CoV than among 15 coronaviruses infecting human and other animals. The protein was also predicted to contain promiscuous MHC-I and MHC-II T-cell epitopes, and linear B-cell epitopes localized in specific locations and functional domains of the protein. By applying reverse vaccinology and machine learning, we predicted potential vaccine targets for effective and safe COVID-19 vaccine development. We then propose that an “Sp/Nsp cocktail vaccine” containing a structural protein(s) (Sp) and a non-structural protein(s) (Nsp) would stimulate effective complementary immune responses.

## Introduction

The emerging Coronavirus Disease 2019 (COVID-19) pandemic poses a massive crisis to global public health. As of March 11, 2020, there were 118,326 confirmed cases and 4,292 deaths, according to the World Health Organization (WHO), and WHO declared the COVID-19 as a pandemic on the same day. As of March 22, there were >300,000 confirmed cases and >10,000 deaths globally in at least 167 countries, and the USA reported >27,000 confirmed cases and >300 deaths. It is critical to develop an effective and safe vaccine(s) to control this fast-spreading disease and stop the pandemic.

The causative agent of the COVID-19 disease is the severe acute respiratory syndrome coronavirus 2 (SARS-CoV-2). Coronaviruses can cause animal diseases such as avian infectious bronchitis caused by the infectious bronchitis virus (IBV), and pig transmissible gastroenteritis caused by a porcine coronavirus^1^. Bats are commonly regarded as the natural reservoir of coronaviruses, which can be transmitted to humans and other animals after genetic mutations. There are seven known human coronaviruses, including the novel SARS-CoV-2. Four of them (HCoV-HKU1, HCoV-OC43, HCoV-229E, and HCoV-NL63) have been circulating in the human population worldwide and cause mild symptoms^2^. Coronavirus became prominence after Severe acute respiratory syndrome (SARS) and Middle East Respiratory Syndrome (MERS) outbreaks. In 2003, the SARS disease caused by the SARS-associated coronavirus (SARS-CoV) infected over 8,000 people worldwide and was contained in the summer of 2003^3^. SARS-CoV-2 and SARS-CoV share high sequence identity^4^. The MERS disease infected more than 2,000 people, which is caused by the MERS-associated coronavirus (MERS-CoV) and was first reported in Saudi Arabia and spread to several other countries since 2012^5^.

Although great efforts have been made to develop and manufacture COVID-19 vaccines, there is no human vaccine on the market to prevent this highly infectious disease. Coronaviruses are positively-stranded RNA viruses with its genome packed inside the nucleocapsid (N) protein and enveloped by the membrane (M) protein, envelope (E) protein, and the spike (S) protein^6^. While many coronavirus vaccine studies targeting different structural proteins were conducted, most of these efforts eventually ceased soon after the outbreak of SARS and MERS. With the recent COVID-19 pandemic outbreak, it is urgent to resume the coronavirus vaccine research. As the immediate response to the on-going pandemic, the first testing in humans of the mRNA-based vaccine targeting the S protein of SARS-CoV-2 (ClinicalTrials.gov Identifier: NCT04283461, Table 1) started on March 16, 2020. As the most superficial and protrusive protein of the coronaviruses, S protein plays a crucial role in mediating virus entry. In the SARS vaccine development, the full-length S protein and its S1 subunit (which contains receptor binding domain) have been frequently used as the vaccine antigens due to their ability to induce neutralizing antibodies that prevent host cell entry and infection.

**Table 1.** Reported SARS-CoV, MERS-CoV, SARS-CoV-2 vaccine clinical trials.

| **Virus** | **Location** | **Phase** | **Year** | **Identifier** | **Vaccine Type** |
| --- | --- | --- | --- | --- | --- |
| SARS-CoV | United States | I | 2004 | NCT00099463 | recombinant DNA vaccine (S protein) |
| SARS-CoV | United States | I | 2007 | NCT00533741 | whole virus vaccine |
| SARS-CoV | United States | I | 2011 | NCT01376765 | recombinant protein vaccine (S protein) |
| MERS | United Kingdom | I | 2018 | NCT03399578 | vector vaccine (S protein) |
| MERS | Germany | I | 2018 | NCT03615911 | vector vaccine (S protein) |
| MERS | Saudi Arabia | I | 2019 | NCT04170829 | vector vaccine (S protein) |
| MERS | Germany, Netherland | I | 2019 | NCT04119440 | vector vaccine (S protein) |
| MERS | Russia | I,II | 2019 | NCT04128059 | vector vaccine (protein not specified) |
| MERS | Russia | I,II | 2019 | NCT04130594 | vector vaccine (protein not specified) |
| SARS-CoV2 | United States | I | 2020 | NCT04283461 | mRNA-based vaccine (S protein) |
| SARS-CoV2 | China | I | 2020 | NCT04313127 | vector vaccine (S protein) |

However, the current coronavirus vaccines, including S protein-based vaccines, might have issues in the lack of inducing complete protection and possible safety concerns^7,8^. All existing SARS/MERS vaccines were reported to induce neutralizing antibodies and partial protection against the viral challenges in animal models (Table 2), but it is desired to induce complete protection or sterile immunity. Moreover, it has become increasingly clear that multiple immune responses, including those induced by humoral or cell-mediated immunity, are responsible for correlates of protection than antibody titers alone^9^. Both killed SARS-CoV whole virus vaccine and adenovirus-based recombinant vector vaccines expressing S or N proteins induced neutralizing antibody responses but did not provide complete protection in animal model^10^. A study has shown increased liver pathology in the vaccinated ferrets immunized with modified vaccinia Ankara-S recombinant vaccine^11^. The safety and efficacy of these vaccination strategies have not been fully tested in human clinical trials, but the safety can be a major concern. Therefore, novel strategies are needed to enhance the efficacy and safety of COVID-19 vaccine development.

**Table 2.**
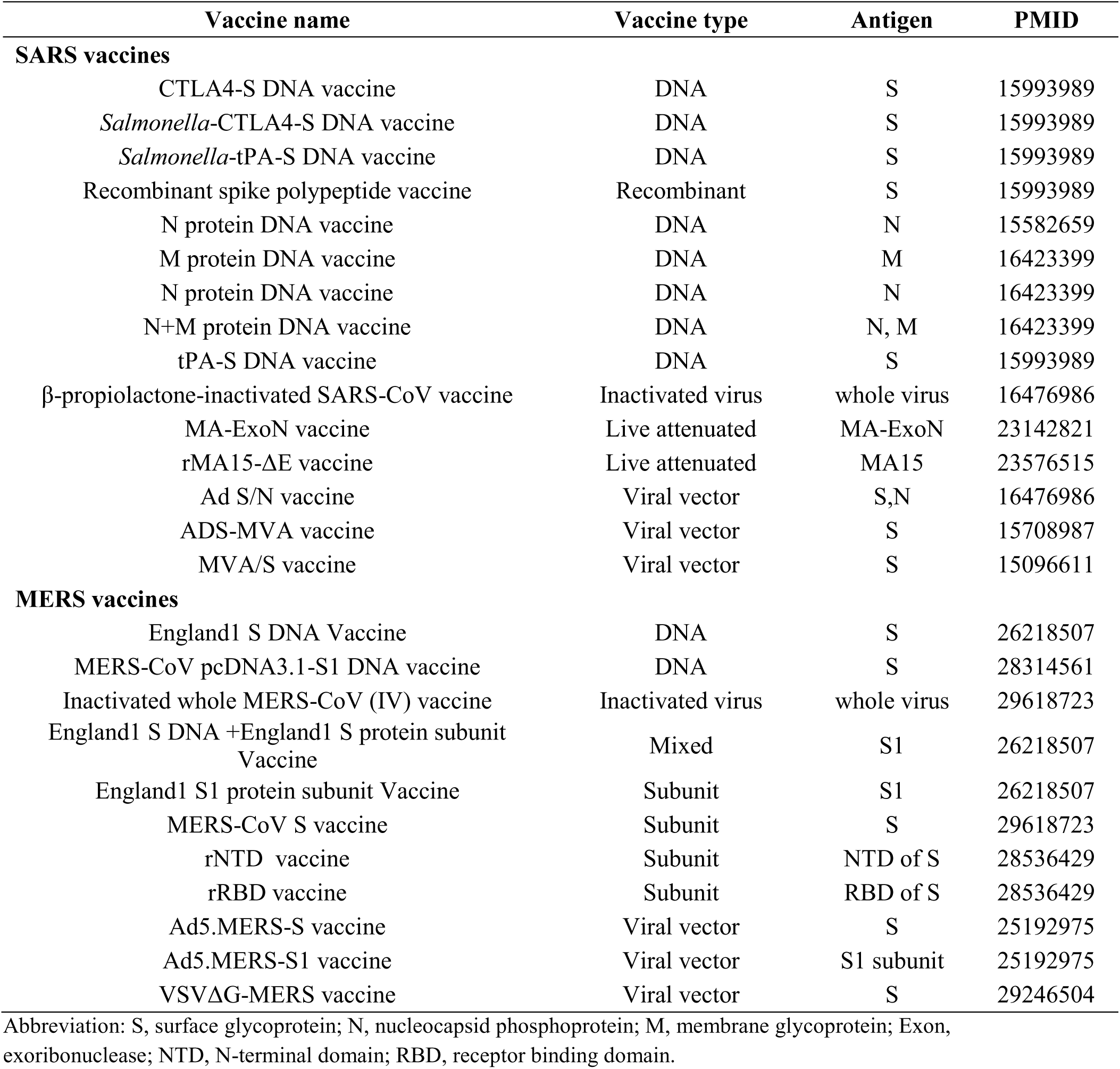
Vaccines tested for SARS-CoV and MERS-CoV.

In recent years, the development of vaccine design has been revolutionized by the reverse vaccinology (RV), which aims to first identify promising vaccine candidate through bioinformatics analysis of the pathogen genome. RV has been successfully applied to vaccine discovery for pathogens such as Group B meningococcus and led to the license Bexsero vaccine^12^. Among current RV prediction tools^13,14^, Vaxign is the first web-based RV program^15^ and has been used to successfully predict vaccine candidates against different bacterial and viral pathogens^16–18^. Recently we have also developed a machine learning approach called Vaxign-ML to enhance prediction accuracy^19^.

In this study, we first surveyed the existing coronavirus vaccine development status, and then applied the Vaxign RV and Vaxign-ML approaches to predict COVID-19 protein candidates for vaccine development. We identified six possible adhesins, including the structural S protein and five other non-structural proteins, and three of them (S, nsp3, and nsp8 proteins) were predicted to induce high protective immunity. The S protein was predicted to have the highest protective antigenicity score, and it has been extensively studied as the target of coronavirus vaccines by other researchers. The sequence conservation and immunogenicity of the multi-domain nsp3 protein, which was predicted to have the second-highest protective antigenicity score yet, was further analyzed in this study. Based on the predicted structural S protein and non-structural proteins (including nsp3) using reverse vaccinology and machine learning, we proposed and discussed a cocktail vaccine strategy, for rational COVID-19 vaccine development.

## Results

### Published research and clinical trial coronavirus vaccine studies

To better understand the current status of coronavirus vaccine development, we systematically surveyed the development of vaccines for coronavirus from the ClinicalTrials.gov database and PubMed literature (as of March 17, 2020). Extensive effort has been made to develop a safe and effective vaccine against SARS or MERS, and the most advance clinical trial study is currently at phase II (Table 1). It is a challenging task to quickly develop a safe and effective vaccine for the on-going COVID-19 pandemic.

There are two primary design strategies for coronavirus vaccine development: the usage of the whole virus or genetically engineered vaccine antigens that can be delivered through different formats. The whole virus vaccines include inactivated^20^ or live attenuated vaccines^21,22^ (Table 2). The two live attenuated SARS vaccines mutated the exoribonuclease and envelop protein to reduce the virulence and/or replication capability of the SARS-CoV. Overall, the whole virus vaccines can induce a strong immune response and protect against coronavirus infections. Genetically engineered vaccines that target specific coronavirus protein are often used to improve vaccine safety and efficacy. The coronavirus antigens such as S protein, N protein, and M protein can be delivered as recombinant DNA vaccine and viral vector vaccine (Table 2).

### N protein is conserved among SARS-CoV-2, SARS-CoV, and MERS-CoV, but missing from the other four human coronaviruses causing mild symptoms

We first used the Vaxign analysis framework^15,19^ to compare the full proteomes of seven human coronavirus strains (SARS-CoV-2, SARS-CoV, MERS-CoV, HCoV-229E, HCoV-OC43, HCoV-NL63, and HCoV-HKU1). The proteins of SARS-CoV-2 were used as the seed for the pan-genomic comparative analysis. The Vaxign pan-genomic analysis reported only the N protein in SARS-CoV-2 having high sequence similarity among the more severe form of coronavirus (SARS-CoV and MERS-CoV), while having low sequence similarity among the more typically mild HCoV-229E, HCoV-OC43, HCoV-NL63, and HCoV-HKU1. The sequence conservation suggested the potential of N protein as a candidate for the cross-protective vaccine against SARS and MERS. The N protein was also evaluated and used for vaccine development (Table 2). The N protein packs the coronavirus RNA to form the helical nucleocapsid in virion assembly. This protein is more conserved than the S protein and was reported to induce an immune response and neutralize coronavirus infections^23^. However, a study also showed the linkage between N protein and severe pneumonia or other serious liver failures related to the pathogenesis of SARS^24^.

### Six adhesive proteins in SARS-CoV-2 identified as potential vaccine targets

The Vaxign RV analysis predicted six SARS-CoV-2 proteins (S protein, nsp3, 3CL-PRO, and nsp8-10) as adhesive proteins (Table 3). Adhesin plays a critical role in the virus adhering to the host cell and facilitating the virus entry to the host cell^25^, which has a significant association with the vaccine-induced protection^26^. In SARS-CoV-2, S protein was predicted to be adhesin, matching its primary role in virus entry. The structure of SARS-CoV-2 S protein was determined^27^ and reported to contribute to the host cell entry by interacting with the angiotensin-converting enzyme 2 (ACE2)^28^. Besides S protein, the other five predicted adhesive proteins were all non-structural proteins. In particular, nsp3 is the largest non-structural protein of SARS-CoV-2 comprises various functional domains^29^.

**Table 3.**
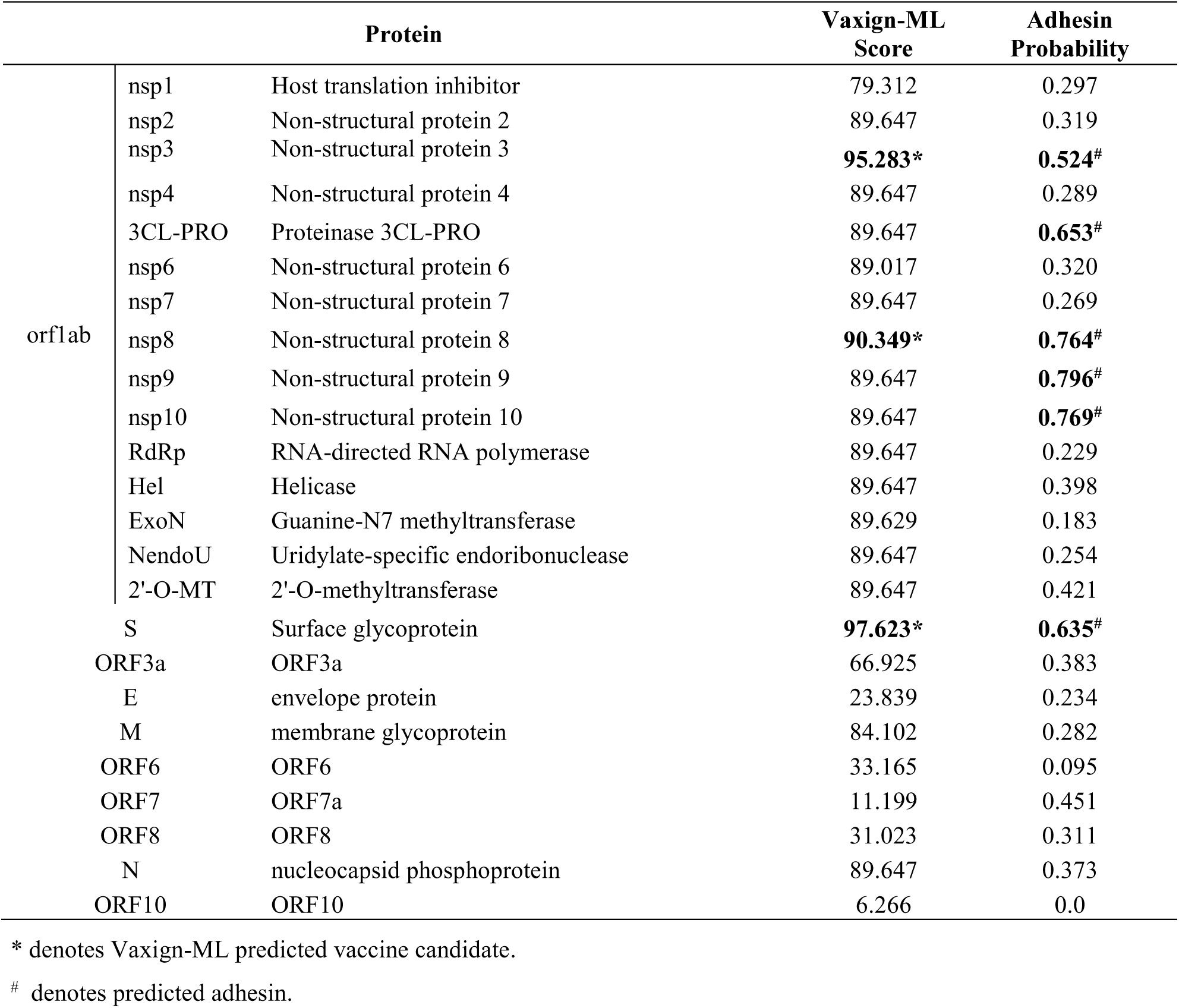
Vaxign-ML Prediction and adhesin probability of all SARS-CoV-2 proteins.

### Three adhesin proteins were predicted to induce strong protective immunity

The Vaxign-ML pipeline computed the protegenicity (protective antigenicity) score and predicted the induction of protective immunity by a vaccine candidate^19^. The training data consisted of viral protective antigens, which were tested to be protective in at least one animal challenge model^30^. The performance of the Vaxign-ML models was evaluated (Table S1 and Figure S1), and the best performing model had a weighted F1-score of 0.94. Using the optimized Vaxign-ML model, we predicted three proteins (S protein, nsp3, and nsp8) as vaccine candidates with significant protegenicity scores (Table 3). The S protein was predicted to have the highest protegenicity score, which is consistent with the experimental observations reported in the literature. The nsp3 protein is the second most promising vaccine candidate besides S protein. There was currently no study of nsp3 as a vaccine target. The structure and functions of this protein have various roles in coronavirus infection, including replication and pathogenesis (immune evasion and virus survival) ^29^. Therefore, we selected nsp3 for further investigation, as described below.

### Nsp3 as a vaccine candidate

The multiple sequence alignment and the resulting phylogeny of nsp3 protein showed that this protein in SARS-CoV-2 was more closely related to the human coronaviruses SARS-CoV and MERS-CoV, and bat coronaviruses BtCoV/HKU3, BtCoV/HKU4, and BtCoV/HKU9. We studied the genetic conservation of nsp3 protein (Figure 1A) in seven human coronaviruses and eight coronaviruses infecting other animals (Table S2). The five human coronaviruses, SARS-CoV-2, SARS-CoV, MERS-CoV, HCoV-HKU1, and HCoV-OC43, belong to the beta-coronavirus while HCoV-229E and HCoV-NL63 belong to the alpha-coronavirus. The HCoV-HKU1 and HCoV-OC43, as the human coronavirus with mild symptoms clustered together with murine MHV-A59. The more severe form of human coronavirus SARS-CoV-2, SARS-CoV, and MERS-CoV grouped with three bat coronaviruses BtCoV/HKU3, BtCoV/HKU4, and BtCoV/HKU9.

**Figure 1.**
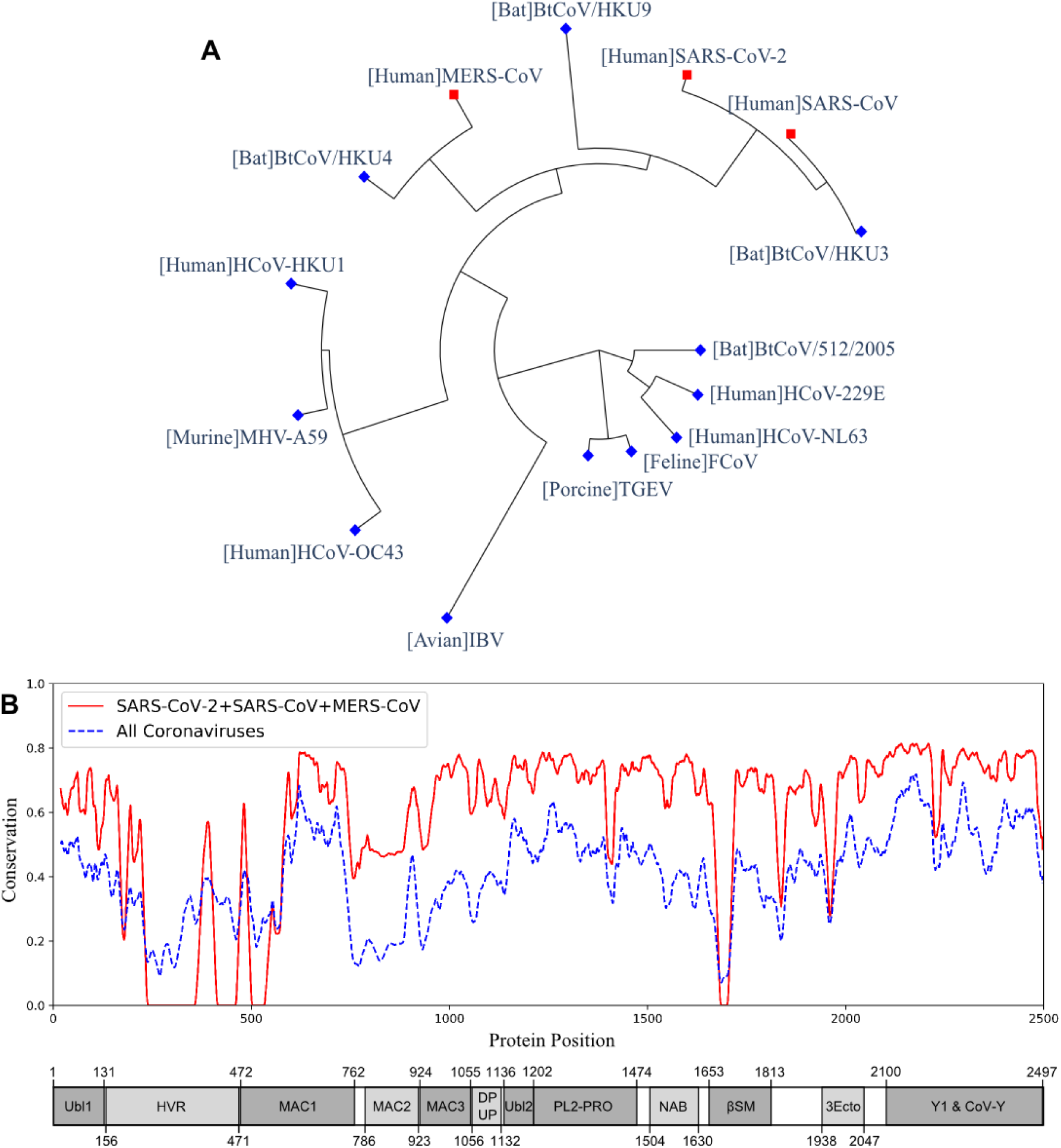
The phylogeny and sequence conservation of coronavirus nsp3. (A) Phylogeny of 15 strains based on the nsp3 protein sequence alignment and phylogeny analysis. (B) The conservation of nsp3 among different coronavirus strains. The red line represents the conservation among the four strains (SARS-CoV, SARS-CoV-2, MERS, and BtCoV-HKU3). The blue line was generated using all the 15 strains. The bottom part represents the nsp3 peptides and their sizes. The phylogenetically close four strains have more conserved nsp3 sequences than all the strains being considered.

When evaluating the amino acid conservations relative to the functional domains in nsp3, all protein domains, except the hypervariable region (HVR), macro-domain 1 (MAC1) and beta-coronavirus-specific marker βSM, showed higher conservation in SARS-CoV-2, SARS-CoV, and MERS-CoV (Figure 1B). The amino acid conservation between the major human coronavirus (SARS-CoV-2, SARS-CoV, and MERS-CoV) was plotted and compared to all 15 coronaviruses used to generate the phylogenetic of nsp3 protein (Figure 1B). The SARS-CoV domains were also plotted (Figure 1B), with the relative position in the multiple sequence alignment (MSA) of all 15 coronaviruses (Table S3 and Figure S2).

The immunogenicity of nsp3 protein in terms of T cell MHC-I & MHC-II and linear B cell epitopes was also investigated. There were 28 and 42 promiscuous epitopes predicted to bind the reference MHC-I & MHC-II alleles, which covered the majority of the world population, respectively (Table S4-5). In terms of linear B cell epitopes, there were 14 epitopes with BepiPred scores over 0.55 and had at least ten amino acids in length (Table S6). The 3D structure of SARS-CoV-2 protein was plotted and highlighted with the T cell MHC-I & MHC-II, and linear B cell epitopes (Figure 2). The predicted B cell epitopes were more likely located in the distal region of the nsp3 protein structure. Most of the predicted MHC-I & MHC-II epitopes were embedded inside the protein. The sliding averages of T cell MHC-I & MHC-II and linear B cell epitopes were plotted with respect to the tentative SARS-CoV-2 nsp3 protein domains using SARS-CoV nsp3 protein as a reference (Figure 3). The ubiquitin-like domain 1 and 2 (Ubl1 and Ubl2) only predicted to have MHC-I epitopes. The Domain Preceding Ubl2 and PL2-PRO (DPUP) domain had only predicted MHC-II epitopes. The PL2-PRO contained both predicted MHC-I and MHC-II epitopes, but not B cell epitopes. In particular, the TM1, TM2, and AH1 were predicted helical regions with high T cell MHC-I and MHC-II epitopes^31^. The TM1 and TM2 are transmembrane regions passing the endoplasmic reticulum (ER) membrane. The HVR, MAC2, MAC3, nucleic-acid binding domain (NAB), βSM, Nsp3 ectodomain; (3Ecto), Y1, and CoV-Y domain contained predicted B cell epitopes. Finally, the Vaxign RV framework also predicted 2 regions (position 251-260 and 329-337) in the MAC1 domain of nsp3 domain having high sequence similarity to the human mono-ADP-ribosyltransferase PARP14 (NP_060024.2).

**Figure 2.**
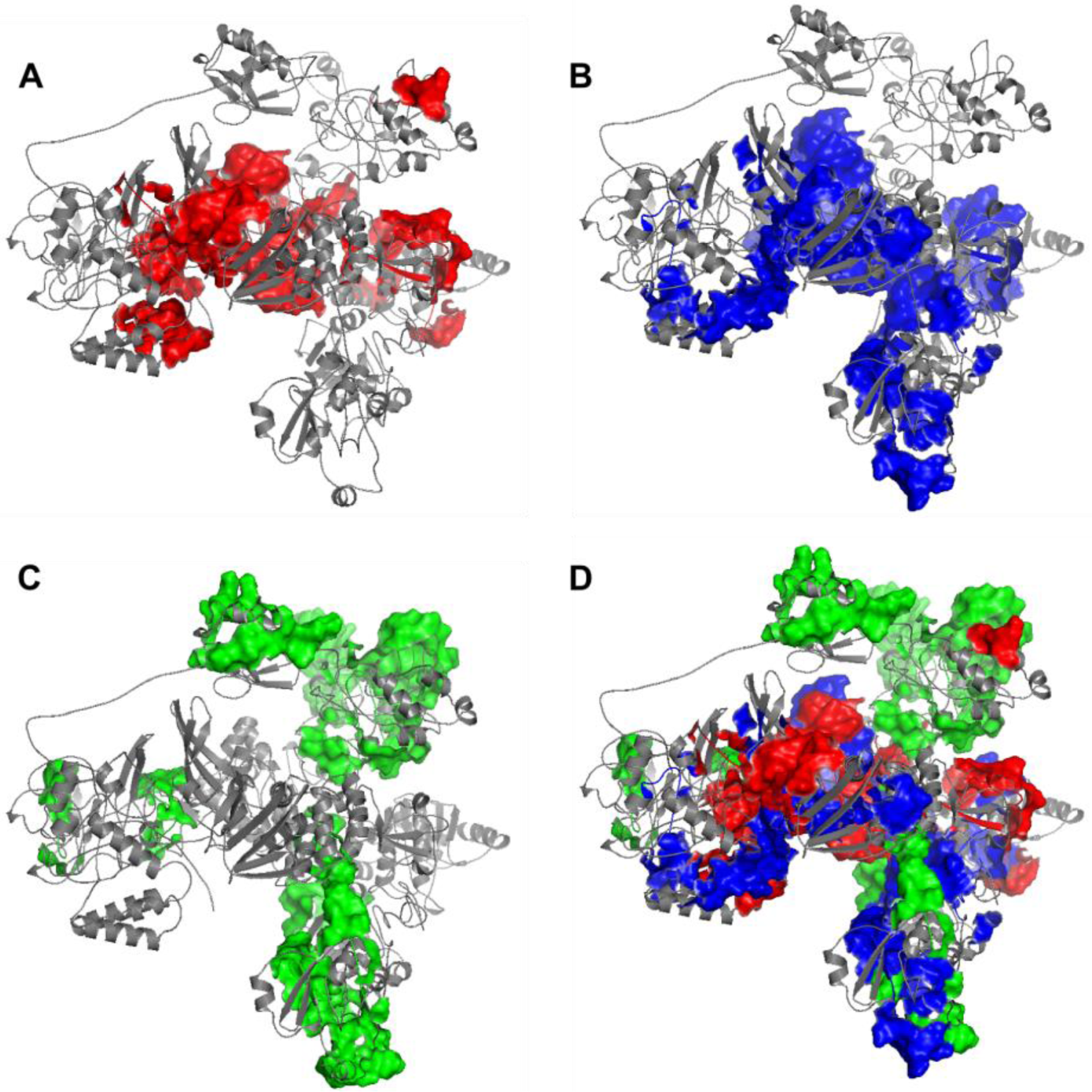
Predicted 3D structure of nsp3 protein highlighted with (A) MHC-I T cell epitopes (red), (B) MHC-II (blue) T cell epitopes, (C) linear B cell epitopes (green), and the merged epitopes. MHC-I epitopes are more internalized, MHC-II epitopes are more mixed, and B cells are more shown on the surface.

**Figure 3.**
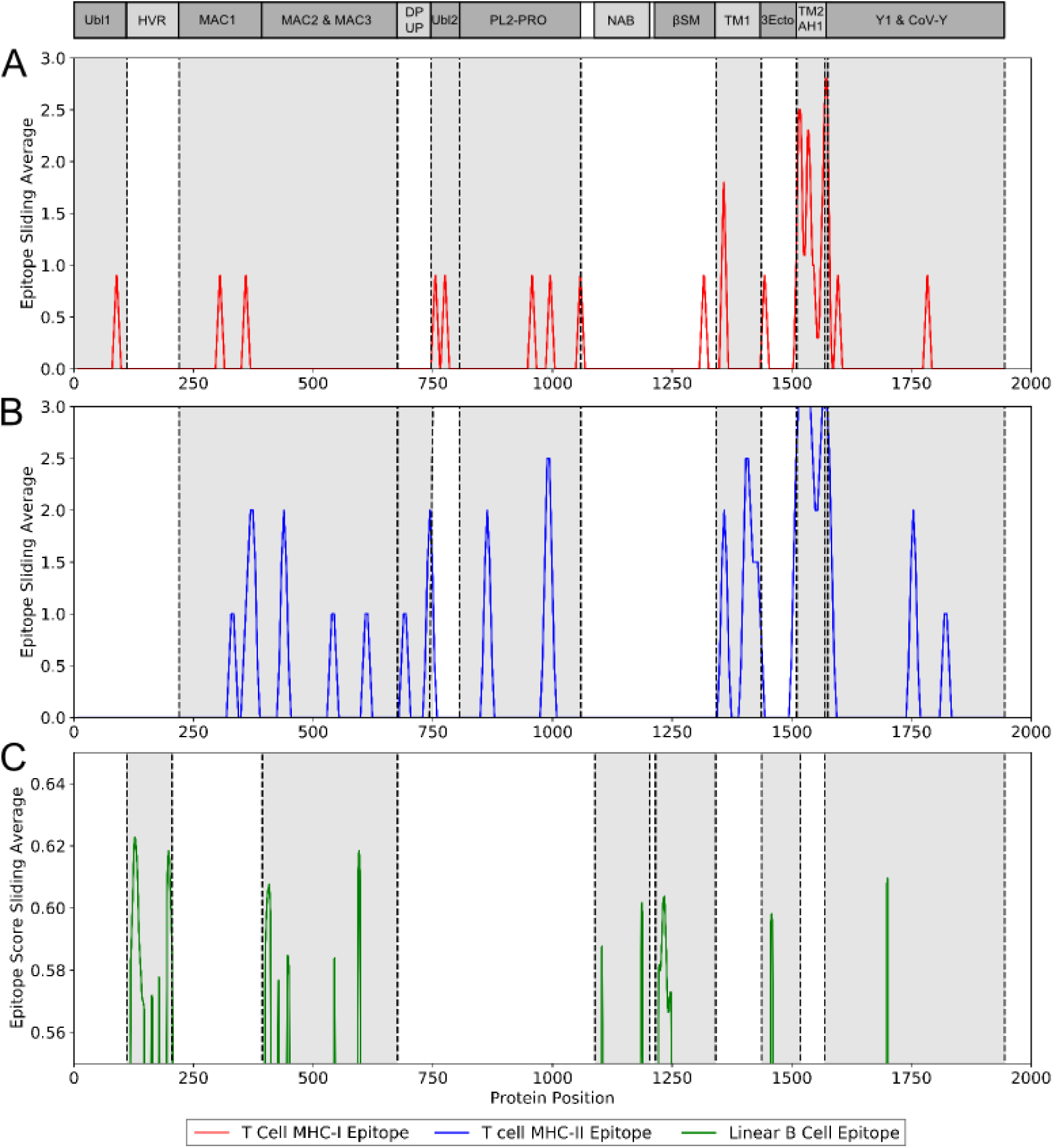
Immunogenic region of nsp3 between SARS-CoV-2 and the four conservation strains. (A) MHC-I (red) T cell epitope (B) MHC-II (blue) T cell epitope (C) linear B cell epitope (green).

## Discussion

Our prediction of the potential SARS-CoV-2 antigens, which could induce protective immunity, provides a timely analysis for the vaccine development against COVID-19. Currently, most coronavirus vaccine studies use the whole inactivated or attenuated virus, or target the structural proteins such as the spike (S) protein, nucleocapsid (N) protein, and membrane (M) protein (Table 2). But the inactivated or attenuated whole virus vaccine might induce strong adverse events. On the other hand, vaccines targeting the structural proteins induce a strong immune response^23,32,33^. In some studies, these structural proteins, including the S and N proteins, were reported to associate with the pathogenesis of coronavirus^24,34^ and might raise safety concern^11^. Our study applied state-of-the-art Vaxign reserve vaccinology (RV) and Vaxign-ML machine learning strategies to the entire SARS-CoV-2 proteomes, including both structural and non-structural proteins for vaccine candidate prediction. Our results indicate, for the first time, that many non-structural proteins could be used as potential vaccine candidates.

The SARS-CoV-2 S protein was identified by our Vaxign and Vaxign-ML analysis as the most favorable vaccine candidate. First, the Vaxign RV framework predicted the S protein as a likely adhesin, which is consistent with the role of S protein for the invasion of host cells. Second, our Vaxign-ML predicted that the S protein had a high protective antigenicity score. These results confirmed the role of S protein as the important target of COVID-19 vaccines. However, targeting only the S protein may induce high serum-neutralizing antibody titers but cannot induce complete protection^10^. In addition, HCoV-NL63 also uses S protein and employs the angiotensin-converting enzyme 2 (ACE2) for cellular entry, despite markedly weak pathogenicity^35^. This suggests that the S protein is not the only factor determining the infection level of a human coronavirus. Thus, alternative vaccine antigens may be considered as potential targets for COVID-19 vaccines.

Among the five non-structural proteins being predicted as potential vaccine candidates, the nsp3 protein was predicted to have second-highest protective antigenicity score, adhesin property, promiscuous MHC-I & MHC-II T cell epitopes, and B cell epitopes. The nsp3 is the largest non-structural protein that includes multiple functional domains related to viral pathogenesis^29^. The multiple sequence alignment of nsp3 also showed higher sequence conservation in most of the functional domains in SARS-CoV-2, SARS-CoV, and MERS-CoV, than in all 15 coronavirus strains (Fig. 1B). Besides the nsp3 protein, our study also predicted four additional non-structural proteins (3CL-pro, nsp8, nsp9, and nsp10) as possible vaccine candidates based on their adhesin probabilities, and the nsp8 protein was also predicted to have a significant protective antigenicity score.

However, these predicted non-structural proteins (nasp3, 3CL-pro, nsp8, nsp9, and nsp10) are not part of the viral structural particle, and all the current SARS/MERS/COVID-19 vaccine studies target the structural (S/M/N) proteins. Although structural proteins are commonly used as viral vaccine candidates, non-structural proteins correlates to vaccine protection. The non-structural protein NS1 was found to induce protective immunity against the infections by flaviviruses^36^. Since NS1 is not part of the virion, antibodies against NS1 have no neutralizing activity but some exhibit complement-fixing activity^37^. However, passive transfer of anti-NS1 antibody or immunization with NS1 conferred protection^38^. Anti-NS1 antibody could also reduce viral replication by complement-dependent cytotoxicity of infected cells, block NS1-induced pathogenic effects, and attenuate NS1-induced disease development during the critical phase^39^. Finally, NS1 is not a structural protein and anti-NS1 antibody will not induce antibody-dependent enhancement (ADE), which is a virulence factor and a risk factor causing many adverse events^39^. The non-structural proteins of the hepatitis C virus were reported to induce HCV-specific vigorous and broad-spectrum T-cell responses^40^. The non-structural HIV-1 gene products were also shown to be valuable targets for prophylactic or therapeutic vaccines^41^. Therefore, it is reasonable to consider the SARS-CoV-2 non-structural proteins (e.g., nsp3) as possible vaccine targets, which might induce cell-mediated or humoral immunity necessary to prevent viral invasion and/or replication. None of the non-structural proteins have been evaluated as vaccine candidates, and the feasibilit of these proteins as vaccine targets are subject to further experimental verification.

In addition to vaccines expressing a single or a combination of structural proteins, here we propose an “Sp/Nsp cocktail vaccine” as an effective strategy for COVID-19 vaccine development. A typical cocktail vaccine includes more than one antigen to cover different aspects of protection^42,43^. The licensed Group B meningococcus Bexsero vaccine, which was developed via reverse vaccinology, contains three protein antigens^12^. To develop an efficient and safe COVID-19 cocktail vaccine, an “Sp/Nsp cocktail vaccine”, which mixes a structural protein(s) (Sp, such as S protein) and a non-structural protein(s) (Nsp, such as nsp3) could induce more favorable protective immune responses than vaccines expressing a structural protein(s). The benefit of a cocktail vaccine strategy could induce immunity that can protect the host against not only the S-ACE2 interaction and viral entry to the host cells, but also protect against the accessary non-structural adhesin proteins (e.g., nsp3), which might also be vital to the viral entry and replication. The usage of more than one antigen allows us to reduce the volume of each antigen and thus to reduce the induction of adverse events. Nonetheless, the potentials of the proposed “Sp/Nsp cocktail vaccine” strategy need to be experimentally validated.

For rational COVID-19 vaccine development, it is critical to understand the fundamental host-coronavirus interaction and protective immune mechanism^7^. Such understanding may not only provide us guidance in terms of antigen selection but also facilitate our design of vaccine formulations. For example, an important foundation of our prediction in this study is based on our understanding of the critical role of adhesin as a virulence factor as well as protective antigen. The choice of DNA vaccine, recombinant vaccine vector, and another method of vaccine formulation is also deeply rooted in our understanding of pathogen-specific immune response induction. Different experimental conditions may also affect results^44,45^. Therefore, it is crucial to understand the underlying molecular and cellular mechanisms for rational vaccine development.

## Methods

### Annotation of literature and database records

We annotated peer-reviewed journal articles stored in the PubMed database and the ClinicalTrials.gov database. From the peer-reviewed articles, we identified and annotated those coronavirus vaccine candidates that were experimentally studied and found to induce protective neutralizing antibody or provided immunity against virulent pathogen challenge.

### Vaxign prediction

The SARS-CoV-2 sequence was obtained from NCBI. All the proteins of six known human coronavirus strains, including SARS-CoV, MERS-CoV, HCoV-229E, HCoV-OC43, HCoV-NL63, and HCoV-HKU1 were extracted from Uniprot proteomes^46^. The full proteomes of these seven coronaviruses were then analyzed using the Vaxign reverse vaccinology pipeline^15,19^. The Vaxign program predicted serval biological features, including adhesin probability^47^, transmembrane helix^48^, orthologous proteins^49^, and protein functionss^15,19^.

### Vaxign-ML prediction

The ML-based RV prediction model was built following a similar methodology described in the Vaxign-ML^19^. Specifically, the positive samples in the training data included 397 bacterial and 178 viral protective antigens (PAgs) recorded in the Protegen database^30^ after removing homologous proteins with over 30% sequence identity. There were 4,979 negative samples extracted from the corresponding pathogens’ Uniprot proteomes^46^ with sequence dis-similarity to the PAgs, as described in previous studies^50–52^. Homologous proteins in the negative samples were also removed. The proteins in the resulting dataset were annotated with biological and physicochemical features. The biological features included adhesin probability^47^, transmembrane helix^48^, and immunogenicity^53^. The physicochemical features included the compositions, transitions and distributions^54^, quasi-sequence-order^55^, Moreau-Broto auto-correlation^56,57^, and Geary auto-correlation^58^ of various physicochemical properties such as charge, hydrophobicity, polarity, and solvent accessibility^59^. Five supervised ML classification algorithms, including logistic regression, support vector machine, k-nearest neighbor, random forest ^60^, and extreme gradient boosting (XGB) ^61^ were trained on the annotated proteins dataset. The performance of these models was evaluated using a nested five-fold cross-validation (N5CV) based on the area under receiver operating characteristic curve, precision, recall, weighted F1-score, and Matthew’s correlation coefficient. The best performing XGB model was selected to predict the protegenicity score of all SARS-CoV-2 isolate Wuhan-Hu-1 (GenBank ID: MN908947.3) proteins, downloaded from NCBI. A protein with protegenicity score over 0.9 is considered as strong vaccine candidate(weighted F1-score > 0.94 in N5CV).

### Phylogenetic analysis

The protein nsp3 was selected for further investigation. The nsp3 proteins of 14 coronaviruses besides SARS-CoV-2 were downloaded from the Uniprot (Table S2). Multiple sequence alignment of these nsp3 proteins was performed using MUSCLE^62^ and visualized via SEAVIEW^63^. The phylogenetic tree was constructed using PhyML^64^, and the amino acid conservation was estimated by the Jensen-Shannon Divergence (JSD)^65^. The JSD score was also used to generate a sequence conservation line using the nsp3 protein sequences from 4 or 13 coronaviruses.

### Immunogenicity analysis

The immunogenicity of the nsp3 protein was evaluated by the prediction of T cell MHC-I and MHC-II, and linear B cell epitopes. For T cell MHC-I epitopes, the IEDB consensus method was used to predicting promiscuous epitopes binding to 4 out of 27 MHC-I reference alleles with consensus percentile ranking less than 1.0 score^53^. For T cell MHC-II epitopes, the IEDB consensus method was used to predicting promiscuous epitopes binding to more than half of the 27 MHC-II reference alleles with consensus percentile ranking less than 10.0. The MHC-I and MHC-II reference alleles covered a wide range of human genetic variation representing the majority of the world population^66,67^. The linear B cell epitopes were predicted using the BepiPred 2.0 with a cutoff of 0.55 score^68^. Linear B cell epitopes with at least ten amino acids were mapped to the predicted 3D structure of SARS-CoV-2 nsp3 protein visualized via PyMol^69^. The predicted count of T cell MHC-I and MHC-II epitopes, and the predicted score of linear B cell epitopes were computed as the sliding averages with a window size of ten amino acids. The nsp3 protein 3D structure was predicted using C-I-Tasser^70^ available in the Zhang Lab webserver (https://zhanglab.ccmb.med.umich.edu/C-I-TASSER/2019-nCov/).

## Acknowledgments

This work has been supported by the NIH-NIAID grant 1R01AI081062.

## Author contributions

EO and YH contributed to the study design. EO, MW, AH collected the data. EO performed bioinformatics analysis. EO, MW, and YH wrote the manuscript. All authors performed result interpretation, and discussed and reviewed the manuscript.

## References

1. Perlman, S. & Netland, J. Coronaviruses post-SARS: Update on replication and pathogenesis. Nature Reviews Microbiology (2009). doi: 10.1038/nrmicro2147

2. Cabeça, T. K., Granato, C. & Bellei, N. Epidemiological and clinical features of human coronavirus infections among different subsets of patients. Influenza Other Respi. Viruses (2013). doi: 10.1111/irv.12101

3. Lu, R. et al. Genomic characterisation and epidemiology of 2019 novel coronavirus: implications for virus origins and receptor binding. Lancet (2020). doi: 10.1016/S0140-6736(20)30251-8

4. Lai, C.-C., Shih, T.-P., Ko, W.-C., Tang, H.-J. & Hsueh, P.-R. Severe acute respiratory syndrome coronavirus 2 (SARS-CoV-2) and coronavirus disease-2019 (COVID-19): The epidemic and the challenges. Int. J. Antimicrob. Agents (2020). doi: 10.1016/j.ijantimicag.2020.105924

5. Chan, J. F. W. et al. Middle East Respiratory syndrome coronavirus: Another zoonotic betacoronavirus causing SARS-like disease. Clin. Microbiol. Rev. (2015). doi: 10.1128/CMR.00102-14

6. Li, F. Structure, Function, and Evolution of Coronavirus Spike Proteins. Annu. Rev. Virol. (2016). doi: 10.1146/annurev-virology-110615-042301

7. Roper, R. L. & Rehm, K. E. SARS vaccines: Where are we? Expert Review of Vaccines (2009). doi: 10.1586/erv.09.43

8. deWit, E., vanDoremalen, N., Falzarano, D. & Munster, V. J. SARS and MERS: recent insights into emerging coronaviruses. Nat. Rev. Microbiol. 14, 523–534 (2016).

9. Plotkin, S. A. Updates on immunologic correlates of vaccine-induced protection. Vaccine 38, 2250–2257 (2020).

10. See, R. H. et al. Severe acute respiratory syndrome vaccine efficacy in ferrets: Whole killed virus and adenovirus-vectored vaccines. J. Gen. Virol. (2008). doi: 10.1099/vir.0.2008/001891-0

11. Weingartl, H. et al. Immunization with Modified Vaccinia Virus Ankara-Based Recombinant Vaccine against Severe Acute Respiratory Syndrome Is Associated with Enhanced Hepatitis in Ferrets. J. Virol. (2004). doi: 10.1128/jvi.78.22.12672-12676.2004

12. Folaranmi, T., Rubin, L., Martin, S. W., Patel, M. & MacNeil, J. R. Use of Serogroup B Meningococcal Vaccines in Persons Aged >/=10 Years at Increased Risk for Serogroup B Meningococcal Disease: Recommendations of the Advisory Committee on Immunization Practices, 2015. MMWR Morb Mortal Wkly Rep 64, 608–612 (2015).

13. He, Y. et al. Emerging vaccine informatics. J. Biomed. Biotechnol. 2010, (2010).

14. Dalsass, M., Brozzi, A., Medini, D. & Rappuoli, R. Comparison of Open-Source Reverse Vaccinology Programs for Bacterial Vaccine Antigen Discovery. Front. Immunol. 10, 1–12 (2019).

15. He, Y., Xiang, Z. & Mobley, H. L. T. Vaxign: The first web-based vaccine design program for reverse vaccinology and applications for vaccine development. J. Biomed. Biotechnol. 2010, (2010).

16. Xiang, Z. A. & He, Y. O. Genome-wide prediction of vaccine targets for human herpes simplex viruses using Vaxign reverse vaccinology Human Herpes Simplex (HSV) Viruses. 14, 1–10 (2013).

17. Singh, R., Garg, N., Shukla, G., Capalash, N. & Sharma, P. Immunoprotective Efficacy of Acinetobacter baumannii Outer Membrane Protein, FilF, Predicted In silico as a Potential Vaccine Candidate. Front. Microbiol. 7, (2016).

18. Navarro-Quiroz, E. et al. Prediction of Epitopes in the Proteome of Helicobacter pylori. Glob. J. Health Sci. 10, 148 (2018).

19. Ong, E. et al. Vaxign-ML: Supervised Machine Learning Reverse Vaccinology Model for Improved Prediction of Bacterial Protective Antigens. Bioinformatics (2020).

20. See, R. H. et al. Comparative evaluation of two severe acute respiratory syndrome (SARS) vaccine candidates in mice challenged with SARS coronavirus. J. Gen. Virol. (2006). doi: 10.1099/vir.0.81579-0

21. Graham, R. L. et al. A live, impaired-fidelity coronavirus vaccine protects in an aged, immunocompromised mouse model of lethal disease. Nat. Med. (2012). doi: 10.1038/nm.2972

22. Fett, C., DeDiego, M. L., Regla-Nava, J. A., Enjuanes, L. & Perlman, S. Complete Protection against Severe Acute Respiratory Syndrome Coronavirus-Mediated Lethal Respiratory Disease in Aged Mice by Immunization with a Mouse-Adapted Virus Lacking E Protein. J. Virol. (2013). doi: 10.1128/jvi.00087-13

23. Zhao, P. et al. Immune responses against SARS-coronavirus nucleocapsid protein induced by DNA vaccine. Virology (2005). doi: 10.1016/j.virol.2004.10.016

24. Yasui, F. et al. Prior Immunization with Severe Acute Respiratory Syndrome (SARS)-Associated Coronavirus (SARS-CoV) Nucleocapsid Protein Causes Severe Pneumonia in Mice Infected with SARS-CoV. J. Immunol. (2008). doi: 10.4049/jimmunol.181.9.6337

25. Ribet, D. & Cossart, P. How bacterial pathogens colonize their hosts and invade deeper tissues. Microbes Infect. 17, 173–183 (2015).

26. Ong, E., Wong, M. U. & He, Y. Identification of New Features from Known Bacterial Protective Vaccine Antigens Enhances Rational Vaccine Design. Front. Immunol. 8, 1–11 (2017).

27. Wrapp, D. et al. Cryo-EM structure of the 2019-nCoV spike in the prefusion conformation. Science (2020). doi: 10.1126/science.abb2507

28. Letko, M., Marzi, A. & Munster, V. Functional assessment of cell entry and receptor usage for SARS-CoV-2 and other lineage B betacoronaviruses. Nat. Microbiol. (2020). doi: 10.1038/s41564-020-0688-y

29. Lei, J., Kusov, Y. & Hilgenfeld, R. Nsp3 of coronaviruses: Structures and functions of a large multi-domain protein. Antiviral Research 149, 58–74 (2018).

30. Yang, B., Sayers, S., Xiang, Z. & He, Y. Protegen: A web-based protective antigen database and analysis system. Nucleic Acids Res. 39, 1073–1078 (2011).

31. Rothbard, J. B. & Taylor, W. R. A sequence pattern common to T cell epitopes. EMBO J. (1988). doi: 10.1002/j.1460-2075.1988.tb02787.x

32. Shi, S. Q. et al. The expression of membrane protein augments the specific responses induced by SARS-CoV nucleocapsid DNA immunization. Mol. Immunol. (2006). doi: 10.1016/j.molimm.2005.11.005

33. Al-Amri, S. S. et al. Immunogenicity of Candidate MERS-CoV DNA Vaccines Based on the Spike Protein. Sci. Rep. (2017). doi: 10.1038/srep44875

34. Glansbeek, H. L. et al. Adverse effects of feline IL-12 during DNA vaccination against feline infectious peritonitis virus. J. Gen. Virol. (2002). doi: 10.1099/0022-1317-83-1-1

35. Hofmann, H. et al. Human coronavirus NL63 employs the severe acute respiratory syndrome coronavirus receptor for cellular entry. Proc. Natl. Acad. Sci. U. S. A. (2005). doi: 10.1073/pnas.0409465102

36. Salat, J. et al. Tick-borne encephalitis virus vaccines contain non-structural protein 1 antigen and may elicit NS1-specific antibody responses in vaccinated individuals. Vaccines (2020). doi: 10.3390/vaccines8010081

37. Schlesinger, J. J., Brandriss, M. W. & Walsh, E. E. Protection against 17D yellow fever encephalitis in mice by passive transfer of monoclonal antibodies to the nonstructural glycoprotein gp48 and by active immunization with gp48. J. Immunol. (1985).

38. Gibson, C. A., Schlesinger, J. J. & Barrett, A. D. T. Prospects for a virus non-structural protein as a subunit vaccine. Vaccine (1988). doi: 10.1016/0264-410X(88)90004-7

39. Chen, H. R., Lai, Y. C. & Yeh, T. M. Dengue virus non-structural protein 1: A pathogenic factor, therapeutic target, and vaccine candidate. Journal of Biomedical Science (2018). doi: 10.1186/s12929-018-0462-0

40. Ip, P. P. et al. Alphavirus-based vaccines encoding nonstructural proteins of hepatitis c virus induce robust and protective T-cell responses. Mol. Ther. (2014). doi: 10.1038/mt.2013.287

41. Cafaro, A. et al. Anti-tat immunity in HIV-1 infection: Effects of naturally occurring and vaccine-induced antibodies against tat on the course of the disease. Vaccines (2019). doi: 10.3390/vaccines7030099

42. Sealy, R. et al. Preclinical and clinical development of a multi-envelope, DNA-virus-protein (D-V-P) HIV-1 vaccine. International Reviews of Immunology (2009). doi: 10.1080/08830180802495605

43. Millet, P. et al. Immunogenicity of the Plasmodium falciparum asexual blood-stage synthetic peptide vaccine SPf66. Am. J. Trop. Med. Hyg. (1993). doi: 10.4269/ajtmh.1993.48.424

44. He, Y. et al. Updates on the web-based VIOLIN vaccine database and analysis system. Nucleic Acids Res. 42, 1124–1132 (2014).

45. Ong, E. et al. VIO: Ontology classification and study of vaccine responses given various experimental and analytical conditions. BMC Bioinformatics (2019). doi: 10.1186/s12859-019-3194-6

46. The UniProt Consortium. The Universal Protein Resource (UniProt). Nucleic Acids Res. 36, D193–7 (2008).

47. Sachdeva, G., Kumar, K., Jain, P. & Ramachandran, S. SPAAN: A software program for prediction of adhesins and adhesin-like proteins using neural networks. Bioinformatics 21, 483–491 (2005).

48. Krogh, A., Larsson, B., vonHeijne, G. & Sonnhammer, E. L. . Predicting transmembrane protein topology with a hidden Markov model: application to complete genomes. J Mol Biol 305, 567–580 (2001).

49. Li, L., Stoeckert, C. J. & Roos, D. S. OrthoMCL: Identification of ortholog groups for eukaryotic genomes. Genome Res. (2003). doi: 10.1101/gr.1224503

50. Bowman, B. N. et al. Improving reverse vaccinology with a machine learning approach. Vaccine 29, 8156–8164 (2011).

51. Doytchinova, I. a & Flower, D. R. VaxiJen: a server for prediction of protective antigens, tumour antigens and subunit vaccines. BMC Bioinformatics 8, 4 (2007).

52. Heinson, A. I. et al. Enhancing the biological relevance of machine learning classifiers for reverse vaccinology. Int. J. Mol. Sci. 18, (2017).

53. Fleri, W. et al. The immune epitope database and analysis resource in epitope discovery and synthetic vaccine design. Front. Immunol. 8, 1–16 (2017).

54. Dubchak, I., Muchnik, I., Holbrook, S. R. & Kim, S. H. Prediction of protein folding class using global description of amino acid sequence. Proc. Natl. Acad. Sci. U. S. A. 92, 8700–8704 (1995).

55. Chou, K.-C. Prediction of Protein Subcellular Locations by Incorporating Quasi-Sequence-Order Effect. Biochem. Biophys. Res. Commun. 278, 477–483 (2000).

56. Lin, Z. & Pan, X. M. Accurate prediction of protein secondary structural content. Protein J. 20, 217–220 (2001).

57. Feng, Z. P. & Zhang, C. T. Prediction of membrane protein types based on the hydrophobic index of amino acids. J. Protein Chem. 19, 269–275 (2000).

58. Sokal, R. R. & Thomson, B. A. Population structure inferred by local spatial autocorrelation: An example from an Amerindian tribal population. Am. J. Phys. Anthropol. 129, 121–131 (2006).

59. Ong, S. A. K., Lin, H. H., Chen, Y. Z., Li, Z. R. & Cao, Z. Efficacy of different protein descriptors in predicting protein functional families. BMC Bioinformatics 8, 1–14 (2007).

60. Pedregosa, F. et al. Scikit-learn: Machine Learning in Python. J. Mach. Learn. Res. 12, 2825–2830 (2012).

61. Chen, T. & Guestrin, C. XGBoost: A scalable tree boosting system. Proc. ACM SIGKDD Int. Conf. Knowl. Discov. Data Min. 13-17-Augu, 785–794 (2016).

62. Edgar, R. C. MUSCLE: Multiple sequence alignment with high accuracy and high throughput. Nucleic Acids Res. (2004). doi: 10.1093/nar/gkh340

63. Gouy, M., Guindon, S. & Gascuel, O. Sea view version 4: A multiplatform graphical user interface for sequence alignment and phylogenetic tree building. Mol. Biol. Evol. (2010). doi: 10.1093/molbev/msp259

64. Lefort, V., Longueville, J. E. & Gascuel, O. SMS: Smart Model Selection in PhyML. Mol. Biol. Evol. (2017). doi: 10.1093/molbev/msx149

65. Capra, J. A. & Singh, M. Predicting functionally important residues from sequence conservation. Bioinformatics (2007). doi: 10.1093/bioinformatics/btm270

66. Greenbaum, J. et al. Functinal classification of class II human leukocyte antigen (HLA) molecules reveals seven different supertypes and a surprising degree of repertoire sharing across supertypes. Immunogenetics 63, 325–335 (2013).

67. Weiskopf, D. et al. Comprehensive analysis of dengue virus-specific responses supports an HLA-linked protective role for CD8+ T cells. Proc. Natl. Acad. Sci. U. S. A. 110, E2046–53 (2013).

68. Jespersen, M. C., Peters, B., Nielsen, M. & Marcatili, P. BepiPred-2.0: Improving sequence-based B-cell epitope prediction using conformational epitopes. Nucleic Acids Res. 45, W24–W29 (2017).

69. Schrödinger, L. The PyMol Molecular Graphics System, Versión 1.8. Thomas Holder (2015). doi: 10.1007/s13398-014-0173-7.2

70. Zheng, W. et al. Deep-learning contact-map guided protein structure prediction in CASP13. Proteins Struct. Funct. Bioinforma. (2019). doi: 10.1002/prot.25792

