## Supplemental materials for "COVID-19 coronavirus vaccine design using reverse vaccinology and machine learning"

F
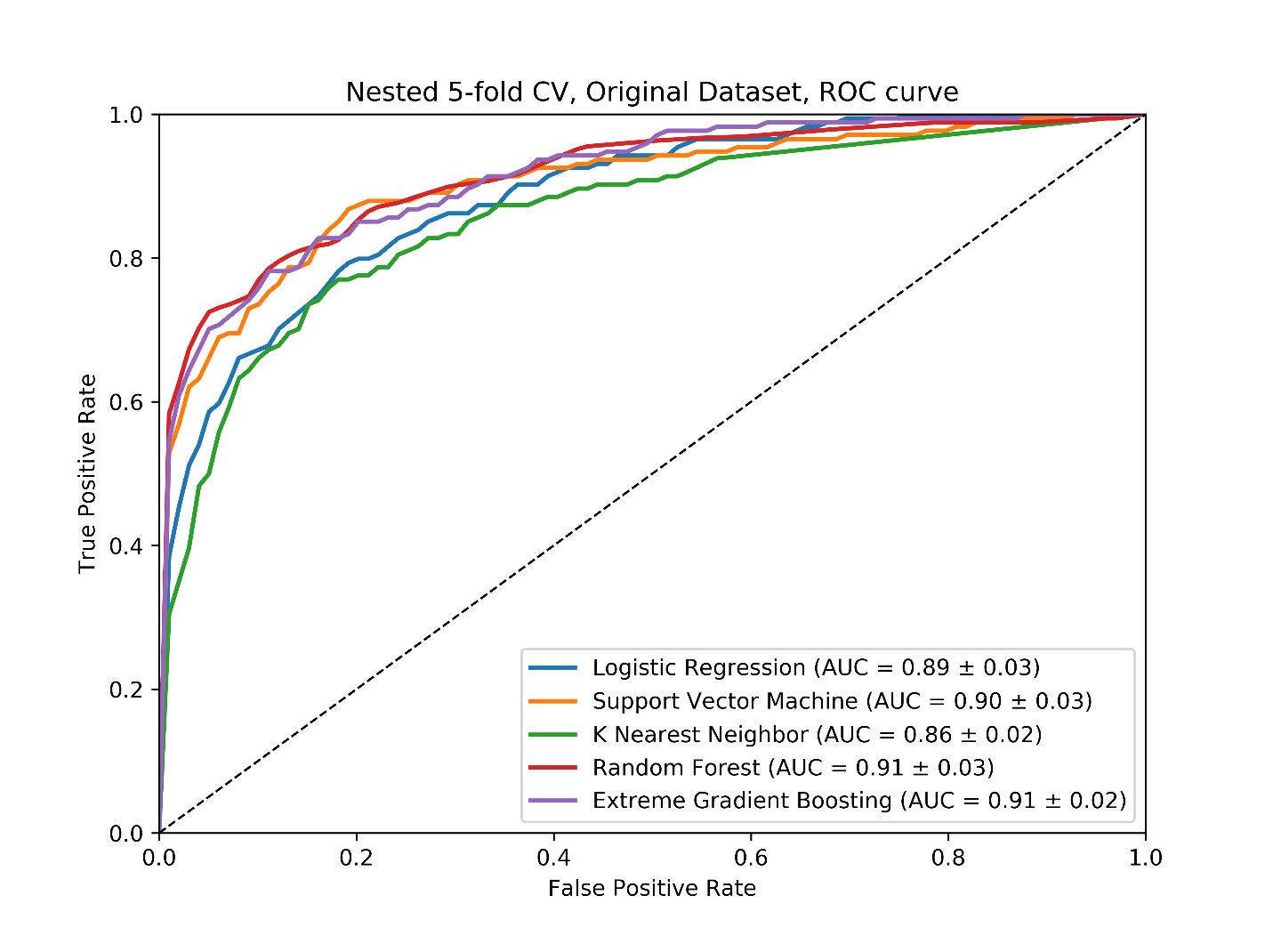

**Supplemental Figure S1.** The average ROC curves in nested five-fold cross validation of five machine learning algorithms (logistic regression, support vector machine, k nearest neighbor, random forest, and extreme gradient boosting).

**Supplemental Figure S2.** Multiple sequence alignment of nsp3 in 15 coronaviruses.

[Please see the attached Figure_S2_nsp3_MSA.]

**Supplemental Table S1.** Nested five-fold cross validation evaluation metrics of five machine learning algorithms.

| **Models** | **Precision** | **Recall** | **Weighted_F1** | **MCC** |
| --- | --- | --- | --- | --- |
| Logistic Regression | 0.541 | 0.366 | 0.886 | 0.370 |
| Support Vector Machine | 0.902 | 0.483 | 0.932 | 0.633 |
| K Nearest Neighbor | 0.489 | 0.552 | 0.895 | 0.458 |
| Random Forest | 0.949 | 0.403 | 0.923 | 0.593 |
| Extreme Gradient Boosting | 0.801 | 0.600 | 0.939 | 0.663 |

**Supplemental Table S2.** The full proteome and nsp3 protein IDs of 15 coronaviruses used in this study.

| **Proteome ID** | **Protein ID** | **Organism** | **Organism Taxon ID** |
| --- | --- | --- | --- |
| UP000000354 | P0C6X7 | Human SARS coronavirus (SARS-CoV) (Severe acute respiratory syndrome coronavirus) | 694009 |
| UP000171868 | T2B9U0 | Middle East respiratory syndrome-related coronavirus | 1335626 |
| UP000006716 | P0C6X1 | Human coronavirus 229E (HCoV-229E) | 11137 |
| UP000001985 | P0C6X4 | Human coronavirus HKU1 (isolate N5) (HCoV-HKU1) (Strain: Isolate N5) | 443241 |
| UP000007552 | P0C6X6 | Human coronavirus OC43 (HCoV-OC43) | 31631 |
| UP000008573 | P0C6X5 | Human coronavirus NL63 (HCoV-NL63) | 277944 |
| UP000000835 | Q98VG9 | Feline coronavirus (strain FIPV WSU-79/1146) (FCoV) (Strain: FIPV WSU-79/1146) | 33734 |
| UP000006717 | P0C6Y1 | Avian infectious bronchitis virus (strain Beaudette) (IBV) (Strain: Beaudette) | 11122 |
| UP000007192 | P0C6X9 | Murine coronavirus (strain A59) (MHV-A59) (Murine hepatitis virus) (Strain: A59) | 11142 |
| UP000001440 | P0C6Y5 | Porcine transmissible gastroenteritis coronavirus (strain Purdue) (TGEV) (Strain: Purdue) | 11151 |
| UP000007451 | P0C6W4 | Bat coronavirus HKU5 (BtCoV) (BtCoV/HKU5/2004) | 694008 |
| UP000006576 | P0C6W5 | Bat coronavirus HKU9 (BtCoV) (BtCoV/HKU9) | 694006 |
| UP000006574 | P0C6W3 | Bat coronavirus HKU4 (BtCoV) (BtCoV/HKU4/2004) (Strain: B04f) | 694007 |
| UP000113079 | P0C6W0 | Bat coronavirus 512/2005 (BtCoV) (BtCoV/512/2005) | 693999 |
| UP000007450 | P0C6W2 | Bat coronavirus HKU3 (BtCoV) (SARS-like coronavirus HKU3) | 442736 |

**Supplemental Table S3.** The position of nsp3 domains in SARS-CoV, and the corresponding relative positive of each domain in the SARS-CoV-2 and the multiple sequence alignment.

|  |  | **nsp3 domains** | | | | | | | | | | | | | | |
| --- | --- | --- | --- | --- | --- | --- | --- | --- | --- | --- | --- | --- | --- | --- | --- | --- |
|  |  | **Ubl1** | **HVR** | **MAC1** | **MAC2** | **MAC3** | **DPUP** | **Ubl2** | **PL2-PRO** | **NAB** | **βSM** | **TM1** | **3Ecto** | **TM2** | **AH1** | **Y1 & CoV-Y** |
| SARS-CoV | start | 1 | 113 | 184 | 389 | 525 | 653 | 723 | 783 | 1066 | 1203 | 1391 | 1414 | 1496 | 1523 | 1546 |
|  | end | 112 | 183 | 365 | 524 | 652 | 720 | 1036 | 1036 | 1180 | 1318 | 1413 | 1495 | 1518 | 1545 | 1922 |
| SARS-CoV-2 | start | 1 | 112 | 206 | 413 | 549 | 677 | 746 | 806 | 1089 | 1226 | 1414 | 1437 | 1519 | 1546 | 1569 |
|  | end | 111 | 205 | 393 | 548 | 676 | 743 | 1059 | 1059 | 1203 | 1341 | 1436 | 1518 | 1541 | 1568 | 1945 |
| MSA relation position | start | 1 | 132 | 472 | 786 | 924 | 1056 | 1136 | 1202 | 1504 | 1653 | 1911 | 1938 | 2048 | 2077 | 2100 |
|  | end | 131 | 471 | 762 | 923 | 1055 | 1132 | 1474 | 1474 | 1630 | 1813 | 1937 | 2047 | 2070 | 2099 | 2497 |

**Supplemental Table S4.** Predicted promiscuous T cell MHC-I epitopes binding to 4 out of 27 reference alleles with consensus percentile ranking over 1.0 using IEDB consensus method.

| **Epitope** | **Start** | **End** | **Allele** |
| --- | --- | --- | --- |
| STNVTIATY | 1455 | 1465 | HLA-A*26:01,HLA-B*15:01,HLA-A*30:02,HLA-A*01:01 |
| RMYIFFASF | 1564 | 1574 | HLA-A*23:01,HLA-A*24:02,HLA-A*32:01,HLA-B*08:01,HLA-B*15:01 |
| AEWFLAYIL | 1507 | 1517 | HLA-B*44:02,HLA-A*32:01,HLA-B*44:03,HLA-B*40:01 |
| MSNLGMPSY | 1436 | 1446 | HLA-A*30:02,HLA-B*35:01,HLA-A*01:01,HLA-B*58:01,HLA-B*15:01 |
| LVAEWFLAY | 1505 | 1515 | HLA-B*35:01,HLA-A*26:01,HLA-B*15:01,HLA-A*01:01 |
| ILFTRFFYV | 1514 | 1524 | HLA-A*02:01,HLA-A*02:06,HLA-A*02:03,HLA-B*08:01 |
| MMSAPPAQY | 988 | 998 | HLA-B*15:01,HLA-A*03:01,HLA-A*30:02,HLA-B*35:01 |
| VMYMGTLSY | 950 | 960 | HLA-A*11:01,HLA-A*03:01,HLA-A*30:02,HLA-A*01:01,HLA-A*32:01,HLA-B*15:01 |
| KENSYTTTI | 1051 | 1061 | HLA-A*32:01,HLA-B*44:03,HLA-B*44:02,HLA-B*40:01 |
| WSMATYYLF | 82 | 92 | HLA-A*23:01,HLA-A*24:02,HLA-B*53:01,HLA-B*58:01,HLA-A*32:01,HLA-B*57:01,HLA-B*15:01 |
| AIMQLFFSY | 1527 | 1537 | HLA-A*11:01,HLA-A*26:01,HLA-A*30:02,HLA-A*32:01,HLA-B*15:01,HLA-B*44:03 |
| FFASFYYVW | 1568 | 1578 | HLA-A*23:01,HLA-A*24:02,HLA-B*53:01,HLA-B*58:01 |
| LAAVNSVPW | 1309 | 1319 | HLA-B*35:01,HLA-B*57:01,HLA-B*58:01,HLA-B*53:01 |
| MPYFFTLLL | 1351 | 1361 | HLA-B*35:01,HLA-B*53:01,HLA-B*51:01,HLA-B*08:01,HLA-B*07:02 |
| LAAIMQLFF | 1525 | 1535 | HLA-B*51:01,HLA-B*35:01,HLA-B*53:01,HLA-B*58:01 |
| STCMMCYKR | 1589 | 1599 | HLA-A*11:01,HLA-A*33:01,HLA-A*31:01,HLA-A*68:01 |
| YIFFASFYY | 1566 | 1576 | HLA-A*11:01,HLA-A*26:01,HLA-A*03:01,HLA-A*30:02,HLA-B*35:01,HLA-A*01:01,HLA-B*15:01,HLA-A*68:01 |
| QMAPISAMV | 1555 | 1565 | HLA-A*68:02,HLA-A*02:01,HLA-A*02:06,HLA-A*02:03 |
| SAMVRMYIF | 1560 | 1570 | HLA-A*32:01,HLA-B*57:01,HLA-B*35:01,HLA-B*08:01 |
| RTNVYLAVF | 352 | 362 | HLA-B*57:01,HLA-B*15:01,HLA-A*32:01,HLA-B*58:01 |
| MSMTYGQQF | 768 | 778 | HLA-B*35:01,HLA-B*53:01,HLA-B*58:01,HLA-B*57:01,HLA-B*15:01 |
| RTIKVFTTV | 748 | 758 | HLA-A*68:02,HLA-A*02:06,HLA-A*32:01,HLA-B*58:01 |
| YMPYFFTLL | 1350 | 1360 | HLA-A*24:02,HLA-A*02:01,HLA-A*02:06,HLA-A*02:03 |
| LAYILFTRF | 1511 | 1521 | HLA-B*35:01,HLA-B*53:01,HLA-B*51:01,HLA-B*58:01,HLA-B*15:01 |
| QLFFSYFAV | 1530 | 1540 | HLA-A*68:02,HLA-A*02:01,HLA-A*02:06,HLA-A*02:03 |
| YVNTFSSTF | 1776 | 1786 | HLA-A*32:01,HLA-A*26:01,HLA-B*15:01,HLA-B*35:01 |
| HFISNSWLM | 1539 | 1549 | HLA-A*23:01,HLA-A*26:01,HLA-B*35:01,HLA-A*24:02 |
| HVVGPNVNK | 298 | 308 | HLA-A*11:01,HLA-A*03:01,HLA-A*30:01,HLA-A*68:01 |

**Supplemental Table S5.** Predicted promiscuous T cell MHC-II epitopes binding at least half of the 27 reference alleles with consensus percentile ranking over 10.0 using IEDB consensus method.

| **Epitope** | **Start** | **End** | **Allele** |
| --- | --- | --- | --- |
| ISNSWLMWLIINLVQ | 1541 | 1557 | HLA-DPA1*03:01,HLA-DRB1*03:01,HLA-DQA1*01:02,HLA-DRB1*04:01,HLA-DQB1*06:02,HLA-DRB1*08:02,HLA-DRB1*15:01,HLA-DQB1*05:01,HLA-DPB1*04:01,HLA-DRB1*11:01,HLA-DPA1*01:03,HLA-DQA1*04:01,HLA-DPB1*04:02,HLA-DRB1*12:01,HLA-DQB1*04:02,HLA-DRB1*04:05,HLA-DPA1*02:01,HLA-DPB1*01:01,HLA-DPA1*01,HLA-DRB5*01:01,HLA-DPB1*02:01,HLA-DRB1*01:01,HLA-DQA1*01:01 |
| LAYILFTRFFYVLGL | 1511 | 1527 | HLA-DPA1*03:01,HLA-DRB1*03:01,HLA-DRB1*08:02,HLA-DRB1*15:01,HLA-DQB1*05:01,HLA-DPB1*04:01,HLA-DRB1*11:01,HLA-DPA1*01:03,HLA-DPB1*14:01,HLA-DPB1*04:02,HLA-DPB1*05:01,HLA-DRB1*12:01,HLA-DRB1*07:01,HLA-DRB3*01:01,HLA-DRB1*04:05,HLA-DPA1*02:01,HLA-DPB1*01:01,HLA-DPA1*01,HLA-DRB5*01:01,HLA-DPB1*02:01,HLA-DRB4*01:01,HLA-DQA1*01:01 |
| AAIMQLFFSYFAVHF | 1526 | 1542 | HLA-DPA1*03:01,HLA-DRB1*04:01,HLA-DRB1*08:02,HLA-DRB1*15:01,HLA-DQB1*05:01,HLA-DPB1*04:01,HLA-DRB1*11:01,HLA-DPA1*01:03,HLA-DPB1*14:01,HLA-DPB1*04:02,HLA-DRB1*12:01,HLA-DPB1*05:01,HLA-DRB1*07:01,HLA-DRB3*01:01,HLA-DRB1*04:05,HLA-DPA1*02:01,HLA-DPB1*01:01,HLA-DPA1*01,HLA-DRB5*01:01,HLA-DPB1*02:01,HLA-DRB4*01:01,HLA-DQA1*01:01 |
| AMVRMYIFFASFYYV | 1561 | 1577 | HLA-DPA1*03:01,HLA-DRB1*04:01,HLA-DRB1*08:02,HLA-DRB1*15:01,HLA-DQB1*05:01,HLA-DPB1*04:01,HLA-DRB1*11:01,HLA-DPA1*01:03,HLA-DPB1*14:01,HLA-DQA1*05:01,HLA-DPB1*04:02,HLA-DRB1*12:01,HLA-DPB1*05:01,HLA-DRB1*09:01,HLA-DRB3*01:01,HLA-DRB1*04:05,HLA-DPA1*02:01,HLA-DPB1*01:01,HLA-DPA1*01,HLA-DQB1*02:01,HLA-DRB5*01:01,HLA-DPB1*02:01,HLA-DQA1*01:01 |
| YIFFASFYYVWKSYV | 1566 | 1582 | HLA-DPB1*04:02,HLA-DRB1*12:01,HLA-DPB1*05:01,HLA-DPA1*03:01,HLA-DRB1*15:01,HLA-DRB1*07:01,HLA-DQB1*05:01,HLA-DRB5*01:01,HLA-DPB1*02:01,HLA-DPB1*04:01,HLA-DPB1*14:01,HLA-DRB3*01:01,HLA-DRB1*11:01,HLA-DPA1*01:03,HLA-DRB1*08:02,HLA-DRB1*04:05,HLA-DQA1*01:01,HLA-DPA1*02:01,HLA-DPB1*01:01,HLA-DPA1*01 |
| EETKFLTENLLLYID | 426 | 442 | HLA-DQA1*05:01,HLA-DPB1*04:02,HLA-DRB1*13:02,HLA-DQB1*02:01,HLA-DPB1*05:01,HLA-DPA1*03:01,HLA-DRB1*12:01,HLA-DRB1*03:01,HLA-DQB1*03:02,HLA-DPB1*02:01,HLA-DPB1*04:01,HLA-DPB1*14:01,HLA-DRB3*01:01,HLA-DRB1*11:01,HLA-DPA1*01:03,HLA-DRB1*04:01,HLA-DQA1*03:01,HLA-DPA1*02:01,HLA-DPB1*01:01,HLA-DPA1*01 |
| MPYFFTLLLQLCTFT | 1351 | 1367 | HLA-DPA1*03:01,HLA-DRB1*04:01,HLA-DRB1*08:02,HLA-DRB1*15:01,HLA-DQB1*05:01,HLA-DPB1*04:01,HLA-DRB1*11:01,HLA-DPA1*01:03,HLA-DPB1*14:01,HLA-DQA1*05:01,HLA-DPB1*04:02,HLA-DRB1*12:01,HLA-DPB1*05:01,HLA-DRB1*04:05,HLA-DPA1*02:01,HLA-DPB1*01:01,HLA-DPA1*01,HLA-DQB1*02:01,HLA-DRB5*01:01,HLA-DPB1*02:01,HLA-DQA1*01:01 |
| IIIWFLLLSVCLGSL | 1411 | 1427 | HLA-DPA1*03:01,HLA-DRB1*03:01,HLA-DRB1*04:01,HLA-DRB1*08:02,HLA-DRB1*15:01,HLA-DQB1*05:01,HLA-DPB1*04:01,HLA-DRB1*11:01,HLA-DPB1*14:01,HLA-DQA1*05:01,HLA-DPB1*04:02,HLA-DRB1*12:01,HLA-DRB1*04:05,HLA-DPA1*02:01,HLA-DPB1*01:01,HLA-DPA1*01,HLA-DQB1*02:01,HLA-DRB5*01:01,HLA-DRB4*01:01,HLA-DRB1*01:01,HLA-DQA1*01:01 |
| VAEWFLAYILFTRFF | 1506 | 1522 | HLA-DQB1*03:02,HLA-DPA1*03:01,HLA-DRB1*04:01,HLA-DRB1*08:02,HLA-DRB1*15:01,HLA-DQB1*05:01,HLA-DPB1*04:01,HLA-DRB1*11:01,HLA-DPA1*01:03,HLA-DPB1*14:01,HLA-DQA1*04:01,HLA-DQA1*05:01,HLA-DPB1*04:02,HLA-DPB1*05:01,HLA-DRB1*12:01,HLA-DRB1*07:01,HLA-DQB1*04:02,HLA-DRB1*04:05,HLA-DQA1*03:01,HLA-DPA1*02:01,HLA-DPB1*01:01,HLA-DPA1*01,HLA-DQB1*02:01,HLA-DRB5*01:01,HLA-DPB1*02:01,HLA-DRB4*01:01,HLA-DQA1*01:01 |
| CKSAFYILPSIISNE | 531 | 547 | HLA-DRB1*12:01,HLA-DRB1*15:01,HLA-DRB1*03:01,HLA-DRB1*07:01,HLA-DQB1*05:01,HLA-DRB1*09:01,HLA-DRB5*01:01,HLA-DPB1*02:01,HLA-DRB1*01:01,HLA-DRB1*11:01,HLA-DRB1*04:01,HLA-DRB1*04:05,HLA-DQA1*01:01,HLA-DPA1*02:01,HLA-DPA1*01:03,HLA-DPB1*14:01,HLA-DRB1*08:02 |
| QQESPFVMMSAPPAQ | 981 | 997 | HLA-DRB3*02:02,HLA-DRB1*13:02,HLA-DRB1*03:01,HLA-DRB1*15:01,HLA-DRB1*09:01,HLA-DRB5*01:01,HLA-DQB1*04:02,HLA-DRB3*01:01,HLA-DQA1*01:02,HLA-DRB1*01:01,HLA-DRB1*04:01,HLA-DRB1*11:01,HLA-DRB1*04:05,HLA-DQB1*06:02,HLA-DRB1*08:02,HLA-DQA1*04:01 |
| LFFSYFAVHFISNSW | 1531 | 1547 | HLA-DPB1*04:02,HLA-DPB1*05:01,HLA-DPA1*03:01,HLA-DRB1*15:01,HLA-DRB1*07:01,HLA-DQB1*05:01,HLA-DRB5*01:01,HLA-DPB1*04:01,HLA-DPB1*02:01,HLA-DPB1*14:01,HLA-DRB3*01:01,HLA-DRB1*11:01,HLA-DPA1*01:03,HLA-DRB1*04:01,HLA-DRB1*04:05,HLA-DQA1*01:01,HLA-DPA1*02:01,HLA-DRB1*08:02,HLA-DPB1*01:01,HLA-DPA1*01 |
| FVMMSAPPAQYELKH | 986 | 1002 | HLA-DRB3*02:02,HLA-DRB1*13:02,HLA-DRB1*12:01,HLA-DRB1*03:01,HLA-DRB1*15:01,HLA-DRB1*09:01,HLA-DRB5*01:01,HLA-DQB1*04:02,HLA-DRB3*01:01,HLA-DRB1*01:01,HLA-DRB1*04:01,HLA-DRB1*11:01,HLA-DRB1*04:05,HLA-DRB1*08:02,HLA-DQA1*04:01 |
| LMWLIINLVQMAPIS | 1546 | 1562 | HLA-DRB1*13:02,HLA-DQB1*03:02,HLA-DPA1*03:01,HLA-DRB1*03:01,HLA-DQA1*01:02,HLA-DRB1*04:01,HLA-DQB1*06:02,HLA-DRB1*08:02,HLA-DRB1*15:01,HLA-DQB1*05:01,HLA-DRB1*11:01,HLA-DPB1*14:01,HLA-DQA1*05:01,HLA-DPB1*04:02,HLA-DRB1*12:01,HLA-DRB1*07:01,HLA-DRB1*04:05,HLA-DQA1*03:01,HLA-DPA1*02:01,HLA-DPB1*01:01,HLA-DQB1*02:01,HLA-DRB5*01:01,HLA-DRB4*01:01,HLA-DRB1*01:01,HLA-DQA1*01:01 |
| LMCQPILLLDQALVS | 1746 | 1762 | HLA-DQA1*05:01,HLA-DPB1*04:02,HLA-DRB1*13:02,HLA-DQB1*02:01,HLA-DRB1*12:01,HLA-DPA1*03:01,HLA-DPA1*01,HLA-DRB1*03:01,HLA-DRB1*15:01,HLA-DPB1*04:01,HLA-DRB3*01:01,HLA-DRB4*01:01,HLA-DRB1*11:01,HLA-DRB1*04:01,HLA-DRB1*01:01,HLA-DRB1*04:05,HLA-DPA1*02:01,HLA-DPB1*14:01,HLA-DRB1*08:02 |
| TAFGLVAEWFLAYIL | 1501 | 1517 | HLA-DQB1*03:02,HLA-DPA1*03:01,HLA-DRB1*15:01,HLA-DQB1*05:01,HLA-DPB1*04:01,HLA-DPA1*01:03,HLA-DPB1*14:01,HLA-DQA1*04:01,HLA-DQA1*05:01,HLA-DPB1*04:02,HLA-DPB1*05:01,HLA-DRB1*07:01,HLA-DQB1*04:02,HLA-DQA1*03:01,HLA-DPA1*02:01,HLA-DPB1*01:01,HLA-DPA1*01,HLA-DQB1*02:01,HLA-DRB5*01:01,HLA-DPB1*02:01,HLA-DQA1*01:01 |
| FAVHFISNSWLMWLI | 1536 | 1552 | HLA-DRB3*02:02,HLA-DPA1*03:01,HLA-DRB1*03:01,HLA-DQA1*01:02,HLA-DRB1*04:01,HLA-DQB1*06:02,HLA-DRB1*08:02,HLA-DRB1*15:01,HLA-DQB1*05:01,HLA-DPB1*04:01,HLA-DRB1*11:01,HLA-DPA1*01:03,HLA-DPB1*14:01,HLA-DPB1*04:02,HLA-DRB1*07:01,HLA-DRB3*01:01,HLA-DRB1*04:05,HLA-DPA1*02:01,HLA-DPB1*01:01,HLA-DPA1*01,HLA-DRB5*01:01,HLA-DPB1*02:01,HLA-DQA1*01:01 |
| SFNYLKSPNFSKLIN | 1396 | 1412 | HLA-DRB3*02:02,HLA-DPB1*05:01,HLA-DRB1*03:01,HLA-DRB1*07:01,HLA-DRB1*15:01,HLA-DRB1*09:01,HLA-DRB5*01:01,HLA-DPB1*04:01,HLA-DRB3*01:01,HLA-DRB1*11:01,HLA-DRB1*04:01,HLA-DRB1*08:02,HLA-DRB1*04:05,HLA-DRB1*01:01,HLA-DPA1*02:01,HLA-DPB1*14:01,HLA-DPA1*01 |
| CYLATALLTLQQIEL | 856 | 872 | HLA-DQB1*03:02,HLA-DPA1*03:01,HLA-DQA1*01:02,HLA-DRB1*04:01,HLA-DQB1*06:02,HLA-DPB1*04:01,HLA-DRB1*11:01,HLA-DPA1*01:03,HLA-DPB1*14:01,HLA-DQA1*04:01,HLA-DQA1*05:01,HLA-DPB1*04:02,HLA-DPB1*05:01,HLA-DQB1*04:02,HLA-DQA1*03:01,HLA-DPA1*02:01,HLA-DPB1*01:01,HLA-DPA1*01,HLA-DQB1*02:01,HLA-DPB1*02:01,HLA-DRB4*01:01 |
| VCTNYMPYFFTLLLQ | 1346 | 1362 | HLA-DPB1*04:02,HLA-DPB1*05:01,HLA-DPA1*03:01,HLA-DRB1*15:01,HLA-DQB1*05:01,HLA-DPB1*04:01,HLA-DPB1*02:01,HLA-DPB1*14:01,HLA-DRB3*01:01,HLA-DPA1*01:03,HLA-DRB1*04:01,HLA-DRB1*04:05,HLA-DQA1*01:01,HLA-DPA1*02:01,HLA-DPB1*01:01,HLA-DPA1*01 |
| DKLVSSFLEMKSEKQ | 366 | 382 | HLA-DPB1*04:02,HLA-DPA1*03:01,HLA-DRB1*03:01,HLA-DRB1*15:01,HLA-DQB1*05:01,HLA-DRB5*01:01,HLA-DPB1*04:01,HLA-DPB1*02:01,HLA-DPB1*14:01,HLA-DRB1*11:01,HLA-DRB1*04:01,HLA-DPA1*01:03,HLA-DRB1*04:05,HLA-DQA1*01:01,HLA-DPA1*02:01,HLA-DPB1*01:01,HLA-DPA1*01 |
| DKNLYDKLVSSFLEM | 361 | 377 | HLA-DRB3*02:02,HLA-DPB1*04:02,HLA-DPB1*05:01,HLA-DPA1*03:01,HLA-DRB1*15:01,HLA-DRB1*07:01,HLA-DRB1*09:01,HLA-DQB1*04:02,HLA-DPB1*04:01,HLA-DPB1*02:01,HLA-DRB1*01:01,HLA-DPA1*01:03,HLA-DRB1*08:02,HLA-DRB1*04:05,HLA-DPA1*02:01,HLA-DPB1*01:01,HLA-DPB1*14:01,HLA-DPA1*01,HLA-DQA1*04:01 |
| VYYSQLMCQPILLLD | 1741 | 1757 | HLA-DPB1*04:02,HLA-DPA1*03:01,HLA-DRB1*03:01,HLA-DQB1*05:01,HLA-DPB1*04:01,HLA-DPB1*02:01,HLA-DRB4*01:01,HLA-DRB1*11:01,HLA-DPA1*01:03,HLA-DRB1*04:01,HLA-DRB1*04:05,HLA-DQA1*01:01,HLA-DPA1*02:01,HLA-DPB1*01:01,HLA-DPB1*14:01,HLA-DPA1*01,HLA-DRB1*01:01 |
| GARFYFYTSKTTVAS | 601 | 617 | HLA-DRB3*02:02,HLA-DPB1*04:02,HLA-DPB1*05:01,HLA-DPA1*03:01,HLA-DPA1*01,HLA-DRB1*15:01,HLA-DRB1*07:01,HLA-DRB1*09:01,HLA-DRB5*01:01,HLA-DPB1*02:01,HLA-DPB1*04:01,HLA-DPB1*14:01,HLA-DRB1*11:01,HLA-DRB1*04:01,HLA-DPA1*01:03,HLA-DRB1*04:05,HLA-DRB1*01:01,HLA-DPA1*02:01,HLA-DPB1*01:01,HLA-DRB1*08:02 |
| FTRFFYVLGLAAIMQ | 1516 | 1532 | HLA-DPA1*03:01,HLA-DRB1*03:01,HLA-DRB1*04:01,HLA-DRB1*08:02,HLA-DRB1*15:01,HLA-DQB1*05:01,HLA-DPB1*04:01,HLA-DRB1*11:01,HLA-DPA1*01:03,HLA-DPB1*14:01,HLA-DQA1*04:01,HLA-DQA1*05:01,HLA-DPB1*04:02,HLA-DRB1*12:01,HLA-DPB1*05:01,HLA-DRB1*07:01,HLA-DRB1*09:01,HLA-DQB1*04:02,HLA-DQB1*03:01,HLA-DRB1*04:05,HLA-DPA1*02:01,HLA-DPB1*01:01,HLA-DPA1*01,HLA-DQB1*02:01,HLA-DRB5*01:01,HLA-DPB1*02:01,HLA-DRB1*01:01,HLA-DQA1*01:01 |
| EVITFDNLKTLLSLR | 731 | 747 | HLA-DRB3*02:02,HLA-DPB1*04:02,HLA-DPB1*05:01,HLA-DPA1*03:01,HLA-DRB1*03:01,HLA-DRB3*01:01,HLA-DRB1*11:01,HLA-DRB1*04:01,HLA-DRB1*01:01,HLA-DRB1*04:05,HLA-DPA1*02:01,HLA-DPB1*01:01,HLA-DPB1*14:01,HLA-DRB1*08:02 |
| FCLEASFNYLKSPNF | 1391 | 1407 | HLA-DRB3*02:02,HLA-DPB1*05:01,HLA-DPA1*01,HLA-DRB1*03:01,HLA-DRB1*07:01,HLA-DRB1*09:01,HLA-DRB5*01:01,HLA-DPB1*02:01,HLA-DPB1*04:01,HLA-DRB1*11:01,HLA-DRB1*04:01,HLA-DPA1*01:03,HLA-DRB1*04:05,HLA-DPA1*02:01,HLA-DPB1*01:01,HLA-DRB1*08:02 |
| MAPISAMVRMYIFFA | 1556 | 1572 | HLA-DPB1*05:01,HLA-DRB1*15:01,HLA-DRB1*03:01,HLA-DRB1*04:05,HLA-DRB1*09:01,HLA-DRB5*01:01,HLA-DPB1*02:01,HLA-DRB3*01:01,HLA-DQA1*01:02,HLA-DRB4*01:01,HLA-DRB1*11:01,HLA-DPA1*01:03,HLA-DRB1*04:01,HLA-DQB1*06:02,HLA-DPA1*02:01,HLA-DRB1*08:02 |
| CLGSLIYSTAALGVL | 1421 | 1437 | HLA-DQA1*05:01,HLA-DRB3*02:02,HLA-DRB1*13:02,HLA-DQB1*02:01,HLA-DRB1*12:01,HLA-DRB1*03:01,HLA-DRB1*07:01,HLA-DRB1*15:01,HLA-DRB1*09:01,HLA-DRB5*01:01,HLA-DPB1*02:01,HLA-DQB1*03:01,HLA-DPB1*14:01,HLA-DRB1*01:01,HLA-DRB1*04:01,HLA-DRB1*11:01,HLA-DRB1*04:05,HLA-DPA1*01:03,HLA-DPA1*02:01,HLA-DPB1*01:01 |
| DNLKTLLSLREVRTI | 736 | 752 | HLA-DPB1*04:02,HLA-DRB1*12:01,HLA-DPA1*03:01,HLA-DRB1*03:01,HLA-DRB1*15:01,HLA-DRB4*01:01,HLA-DRB1*11:01,HLA-DRB1*04:01,HLA-DRB1*01:01,HLA-DRB1*04:05,HLA-DPA1*02:01,HLA-DPB1*01:01,HLA-DPB1*14:01,HLA-DRB1*08:02 |
| INLVQMAPISAMVRM | 1551 | 1567 | HLA-DRB3*02:02,HLA-DQA1*05:01,HLA-DRB1*13:02,HLA-DRB1*12:01,HLA-DRB1*03:01,HLA-DRB1*15:01,HLA-DRB1*07:01,HLA-DRB1*09:01,HLA-DRB5*01:01,HLA-DQB1*04:02,HLA-DQB1*03:01,HLA-DRB4*01:01,HLA-DQA1*01:02,HLA-DRB1*01:01,HLA-DRB1*11:01,HLA-DRB1*04:01,HLA-DRB1*04:05,HLA-DQB1*06:02,HLA-DRB1*08:02,HLA-DQA1*04:01 |
| FKWDLTAFGLVAEWF | 1496 | 1512 | HLA-DQA1*05:01,HLA-DQB1*02:01,HLA-DQB1*03:02,HLA-DPB1*05:01,HLA-DRB1*03:01,HLA-DQB1*05:01,HLA-DRB1*09:01,HLA-DQB1*04:02,HLA-DPB1*04:01,HLA-DRB3*01:01,HLA-DQA1*01:01,HLA-DRB1*04:01,HLA-DRB1*04:05,HLA-DQA1*03:01,HLA-DPA1*02:01,HLA-DPB1*01:01,HLA-DPA1*01,HLA-DQA1*04:01 |
| LTENLLLYIDINGNL | 431 | 447 | HLA-DPB1*04:02,HLA-DRB1*13:02,HLA-DRB1*12:01,HLA-DQB1*03:02,HLA-DPA1*03:01,HLA-DRB1*03:01,HLA-DRB1*15:01,HLA-DQB1*05:01,HLA-DQA1*03:01,HLA-DRB3*01:01,HLA-DRB1*04:01,HLA-DRB1*04:05,HLA-DQA1*01:01,HLA-DPA1*02:01,HLA-DPB1*01:01 |
| YVLGLAAIMQLFFSY | 1521 | 1537 | HLA-DRB1*03:01,HLA-DQA1*01:02,HLA-DRB1*04:01,HLA-DQB1*06:02,HLA-DRB1*08:02,HLA-DRB1*15:01,HLA-DPB1*04:01,HLA-DRB1*11:01,HLA-DPA1*01:03,HLA-DPB1*14:01,HLA-DQA1*04:01,HLA-DQA1*05:01,HLA-DPB1*05:01,HLA-DRB1*12:01,HLA-DRB1*09:01,HLA-DQB1*04:02,HLA-DRB3*01:01,HLA-DQB1*03:01,HLA-DRB1*04:05,HLA-DPA1*02:01,HLA-DPA1*01,HLA-DQB1*02:01,HLA-DRB5*01:01,HLA-DPB1*02:01,HLA-DRB4*01:01 |
| WADNNCYLATALLTL | 851 | 867 | HLA-DQA1*05:01,HLA-DPB1*04:02,HLA-DQB1*02:01,HLA-DPB1*05:01,HLA-DPA1*03:01,HLA-DRB1*07:01,HLA-DPB1*02:01,HLA-DPB1*04:01,HLA-DQA1*01:02,HLA-DRB1*11:01,HLA-DPA1*01:03,HLA-DRB1*04:01,HLA-DQB1*06:02,HLA-DPA1*02:01,HLA-DPB1*01:01,HLA-DPB1*14:01,HLA-DPA1*01 |
| NQHEVLLAPLLSAGI | 321 | 337 | HLA-DPB1*04:02,HLA-DRB1*12:01,HLA-DPA1*03:01,HLA-DRB1*15:01,HLA-DRB1*03:01,HLA-DRB1*09:01,HLA-DPB1*02:01,HLA-DPB1*14:01,HLA-DRB1*11:01,HLA-DRB1*04:01,HLA-DRB1*01:01,HLA-DRB1*04:05,HLA-DPA1*01:03,HLA-DPA1*02:01,HLA-DPB1*01:01,HLA-DRB1*08:02 |
| VLSTFISAARQGFVD | 1811 | 1827 | HLA-DRB3*02:02,HLA-DQB1*03:02,HLA-DRB1*03:01,HLA-DRB1*07:01,HLA-DRB1*09:01,HLA-DRB5*01:01,HLA-DQB1*04:02,HLA-DRB3*01:01,HLA-DQA1*01:02,HLA-DRB1*11:01,HLA-DRB1*04:01,HLA-DQB1*06:02,HLA-DQA1*03:01,HLA-DRB1*08:02,HLA-DQA1*04:01 |
| SFYYVWKSYVHVVDG | 1571 | 1587 | HLA-DPB1*04:02,HLA-DPA1*03:01,HLA-DRB1*15:01,HLA-DRB1*03:01,HLA-DRB1*07:01,HLA-DRB1*09:01,HLA-DRB5*01:01,HLA-DPB1*04:01,HLA-DPB1*02:01,HLA-DRB3*01:01,HLA-DRB1*11:01,HLA-DPA1*01:03,HLA-DRB1*08:02,HLA-DRB1*04:05,HLA-DPA1*02:01,HLA-DPB1*01:01,HLA-DPB1*14:01,HLA-DPA1*01 |
| TKYLVQQESPFVMMS | 976 | 992 | HLA-DRB3*02:02,HLA-DQA1*05:01,HLA-DRB1*13:02,HLA-DQB1*02:01,HLA-DRB1*12:01,HLA-DRB1*03:01,HLA-DRB1*15:01,HLA-DRB1*04:05,HLA-DRB1*09:01,HLA-DRB5*01:01,HLA-DRB3*01:01,HLA-DRB4*01:01,HLA-DQA1*01:02,HLA-DRB1*11:01,HLA-DRB1*04:01,HLA-DRB1*01:01,HLA-DQB1*06:02,HLA-DPA1*02:01,HLA-DPB1*01:01,HLA-DPB1*14:01 |
| EHFIETISLAGSYKD | 681 | 697 | HLA-DPB1*04:02,HLA-DRB1*12:01,HLA-DPA1*03:01,HLA-DRB1*03:01,HLA-DRB1*07:01,HLA-DRB1*04:05,HLA-DRB5*01:01,HLA-DPB1*02:01,HLA-DQA1*01:02,HLA-DRB1*11:01,HLA-DRB1*04:01,HLA-DPA1*01:03,HLA-DQB1*06:02,HLA-DPA1*02:01,HLA-DPB1*14:01,HLA-DRB1*08:02 |
| KSPNFSKLINIIIWF | 1401 | 1417 | HLA-DPB1*04:02,HLA-DRB1*12:01,HLA-DPA1*03:01,HLA-DPB1*05:01,HLA-DRB1*15:01,HLA-DRB1*07:01,HLA-DRB5*01:01,HLA-DRB3*01:01,HLA-DRB4*01:01,HLA-DRB1*11:01,HLA-DRB1*04:01,HLA-DRB1*04:05,HLA-DPA1*02:01,HLA-DPB1*01:01 |
| VRTNVYLAVFDKNLY | 351 | 367 | HLA-DRB1*13:02,HLA-DPB1*05:01,HLA-DRB1*03:01,HLA-DRB1*07:01,HLA-DQB1*05:01,HLA-DRB5*01:01,HLA-DPB1*02:01,HLA-DPB1*04:01,HLA-DPB1*14:01,HLA-DRB1*11:01,HLA-DPA1*01:03,HLA-DQA1*01:01,HLA-DPA1*02:01,HLA-DPB1*01:01,HLA-DPA1*01 |

**Supplemental Table S6.** Predicted linear B cell epitopes using BepiPred 2.0 with a cutoff of 0.55 and at least 10 amino acids.

| **No.** | **Start** | **End** | **Peptide** | **Length** |
| --- | --- | --- | --- | --- |
| 1 | 111 | 148 | EDEEEGDCEEEEFEPSTQYEYGTEDDYQGKPLEFGATS | 38 |
| 2 | 154 | 165 | EEEQEEDWLDDD | 12 |
| 3 | 170 | 180 | VGQQDGSEDNQ | 11 |
| 4 | 187 | 207 | IVEVQPQLEMELTPVVQTIEV | 21 |
| 7 | 392 | 412 | EVKPFITESKPSVEQRKQDDK | 21 |
| 8 | 419 | 429 | EEVTTTLEETK | 11 |
| 9 | 438 | 451 | YIDINGNLHPDSAT | 14 |
| 14 | 536 | 546 | YILPSIISNEK | 11 |
| 15 | 586 | 599 | RKYKGIKIQEGVVD | 14 |
| 28 | 1095 | 1105 | DLVPNQPYPNA | 11 |
| 31 | 1178 | 1189 | NATNKATYKPNT | 12 |
| 32 | 1214 | 1249 | DAQGMDNLACEDLKPVSEEVVENPTIQKDVLECNVK | 36 |
| 34 | 1448 | 1461 | YREGYLNSTNVTIA | 14 |
| 36 | 1691 | 1701 | GQKTYERHSLS | 11 |
