## Supplemental Figure S2 for "COVID-19 coronavirus vaccine design using reverse vaccinology and machine learning"

Alignment: /data/PythonWorkspace/Vaxign-ML_Virus/PL-PROs.all.aln
Seaview [blocks=10 fontsize=10 LETTER-landscape] on Sat Mar 14 20:15:19 2020

```
                    1
[Feline]FCoV        -GDKSVSFSD NVNVREIEP- -VTR-VRLEF EFDNEVVTQV L-EKVIGTKY KFIGTTWEEF EDS----ISE KLDKIFDTLA EQ----GVEL
[Porcine]TGEV       -GDKTVSFSE EVDVQEIAP- -VTR-VKLEF EFDNEIVTGV L-ERAIGTRY KFTGTTWEEF EES----ISE ELDAIFDTLA NQ----GVEL
[Human]HCoV-HKU1    --GRKVNFNE KPVVMEIPS- -LMT-VKVMF DLD-STFDDI L-GKVCSEFE VEKGVTVDDF VAV----VCD AIENALNSCK EHPVV-GYQV
[Bat]BtCoV/HKU4     MPSKKVTFGD V-NTVEVTA- -YRS-VSITY DIH-PVLDAL LSSSKLATFT VEKDLLVEDF VDV----IKD EVLTLLTPLL RGYDIDGFDV
[Human]SARS-CoV     APIKGVTFGE D-TVWEVQG- -YKN-VRITF ELD-ERVDKV L-NEKCSVYT VESGTEVTEF ACV----VAE AVVKTLQPVS DLLTNMGIDL
[Human]SARS-CoV-2   -APTKVTFGD D-TVIEVQG- -YKS-VNITF ELD-ERIDKV L-NEKCSAYT VELGTEVNEF ACV----VAD AVIKTLQPVS ELLTPLGIDL
[Human]MERS-CoV     APVKKVAFGG D-QVHEVAA- -VRS-VTVEY NIH-AVLDTL LASSSLRTFV VDKSLSIEEF ADV----VKE QVSDLLVKLL RGMPIPDFDL
[Bat]BtCoV/HKU9     -GVSKVTFGD E-EVHTIPN- -TVT-VNFSY DVC-EGLDAI L-DKVMAPFQ VEEGTKLEDL ACVVQKAVYE RLSDLFSDCP AELRP--INL
[Human]HCoV-229E    ---GKVSFSD DVEVKDIEP- -VYR-VKLCF EFEDEKLVDV C-EKAIGKKI KHEG-DWDSF CKT----IQS ALSVV--SC- ------YVNL
[Avian]IBV          --GKTVTFGE T-TVQEIPPP DVVP-IKVSI ECCGEPWNTI FKKAYKEPIE VDTDLTVEQL LSV----IYE KMCDDLKLFP E-----APEP
[Murine]MHV-A59     --GKKVEFND KPKVRKIPS- -TRK-IKITF ALD-ATFDSV L-SKACSEFE VDKDVTLDEL LDV----VLD AVESTLSPCK EHDVI-GTKV
[Bat]BtCoV/HKU3     APVKGVTFGE D-TVLEVQG- -YKN-VKITF ELD-VRVDKV L-NEKCSVYT VESGTEVTEF ACV----VAE AVVKTLQPVS DLLTPMGIDL
[Bat]BtCoV/512/2005 ---GVVTISD EVQVRTIDP- -VYK-VRLEY EFEDETLVKV C-EKAIGTKL KVTG-DWSNL LET----LEK AMDVVRQ--- ------HLDV
[Human]HCoV-OC43    --GRRVTFKE QPTVKEIIS- -MPKIIKVFY ELD-NDFNTI L-NTACGVFE VDDTVDMEEF YAV----VID AIEEKLSPCK ELEGV-GAKV
[Human]HCoV-NL63    ---GKISFSD DVIVHDVEP- -THK-VKLIF EFEDDVVTSL C-KKSFGKSI IYTG-DWEGL HEV----LTS AMNVIGQ--- ------HIKL

                   91
[Feline]FCoV        EGY------- --FIYDTCG- --GFDINNPD GVMISQYDLN TAADDKSDSD ASVEDISLIS DNEDVEQIEE DNTSTDDAED VSSVEGETVS
[Porcine]TGEV       EGY------- --FIYDTCG- --GFDIKNPD GIMISQYDIN ITADEKSEVS ASSEEEEVES VEEDPENEIV EA-------S EGAEGTSSQE
[Human]HCoV-HKU1    RAFLNKLNEN VVYLFDEAG- --DEAMASR- -MYCTFAIED VEDVISSEAV ----EDTIDG VVEDTINDDE DVVTGDNDDE DVVTGDNDDE
[Bat]BtCoV/HKU4     EDFIDVPC-- --YVYNQDG- --DCAWSSN- -MTFSINPVE D--------- ----VEEVEE FIEDDYLSDE LP-------- ------IADD
[Human]SARS-CoV     DEWSVATF-- --YLFDDAG- --EENFSSR- -MYCSFYPPD EEEEDDAECE ----EEEIDE TCEHEYGTED DY-------- -------QGL
[Human]SARS-CoV-2   DEWSMATY-- --YLFDESG- --EFKLASH- -MYCSFYPPD EDEEEGDCE- ----EEEFEP STQYEYGTED DY-------- -------QGK
[Human]MERS-CoV     DDFIDAPC-- --YCFNAEG- --DASWSST- -MIFSLHPVE CDEECSEVEA SDLEEGESEC ISETSTEQVD VS-------- ------HEVS
[Bat]BtCoV/HKU9     EDFLTSEC-- --FVYSKDY- --EKILMPE- -MYFSLEDA- ---------- ----VPVDDE MVDDIEDTVE QA-------- ----------
[Human]HCoV-229E    PTY------- --YIYDEEG- --GNDLSLP- -VMISEWPLS VQQAQQ---- ----EATLPD IAEDVVDQVE EV-------- ------NSIF
[Avian]IBV          PPFENV---- --ALVDKNGK DLDCIKSCH- -LIYRDYESD DDI------- ----EEEDAE ECDTDSGEAE EC-------- -----DTNSE
[Murine]MHV-A59     CALLDRLAGD YVYLFDEGG- --DEVIAPR- -MYCSFSAPD DEDCVAADVV DA--DENQDD DAEDSAVLVA DTQEEDGVAK GQVEADSEIC
[Bat]BtCoV/HKU3     DEWSVATF-- --YLFDDAG- --EEKLSSR- -MYCSFYPPD EEEDCEEC-- -----EDEEE TCEHEYGTED DY-------- --------KGL
[Bat]BtCoV/512/2005 PDY------- --FVYDEEG- --GTDLNLT- -IMVSQWPLS SDSEDDFKAV D---DEPNAN TDETVDTFAE DV-------- ---AETQNVQ
[Human]HCoV-OC43    SAFLQKLEDN PLFLFDEAG- --EEVFAPK- -LYCAFTAPE DDDF------ ----LEESDV EEDDVEGEET DL-------- ---TITSAGQ
[Human]HCoV-NL63    PQF------- --YIYDEEG- --GYDVSKP- -VMISQWPIS NDSNGCVVEA STD-FHQLEC IVDDSVREEV DI-------- ------IEQP

                  181
[Feline]FCoV        VVDVEDFVEQ VSLVEENNVL TPAVNPDEQL SSVEKKDE-- ---------- ---------- ---------- -------VSA KNDPWAAAVD
[Porcine]TGEV       EVETVEVADI TSTEEDVDIV EVSAKDDPWA AAVDVQEA-- ---------- ---------- ---------- ---------- ----------
[Human]HCoV-HKU1    DVVTGDNDDE DVVTGD---- --NDDEDVVT GDNDDEDVVT GDNDDEDVVT GDNDDEDVVT GDNNDEDVVT GDNNDEESVT GDNDDQIVVT
[Bat]BtCoV/HKU4     EEAWARAVEE VMPLDDILVA EIELEEDPPL ETALESVE-- ---------- ---------- ---------- ---------- ----------
[Human]SARS-CoV     PLEFGASAET VRVE------ --EEEEEDWL DDTTEQSE-- ---------- ---------- ---------- ---------- ----------
[Human]SARS-CoV-2   PLEFGATSAA LQPE------ --EEQEEDWL DDDSQQTV-- ---------- ---------- ---------- ---------- ----------
[Human]MERS-CoV     DDEWAAAVDE AFPLDE---- --AEDVTESV QEEAQPVE-- ---------- ---------- ---------- ---------- ----------
[Bat]BtCoV/HKU9     ---------- ---------- --SDSDDQWL GDEGAEDC-- ---------- ---------- ---------- ---------- ----------
[Human]HCoV-229E    DIETVDVKHD VSPFEM---- --PFEELNGL KILKQLDN-- ---------- ---------- ---------- ---------- ----------
[Avian]IBV          CEEEDEDTKV LALIQD---- --PASIKYPL PLDEDYSV-- ---------- ---------- ---YNGCIVH ---------- ----------
[Murine]MHV-A59     VAHTGSQEEL AEPDAVGSQT PIASAEETEV GEASDREGIA EAKATVCADA VDACPDQVEA FEIEKVEDSI ---------- ----------
[Bat]BtCoV/HKU3     PLEFGASTET PHVEE----- --EEEEEDWL DDAIEAEP-- ---------- ---------- ---------- ---------- ----------
[Bat]BtCoV/512/2005 QDVTQDEVEA VCDLVV---- --KATEEGPI EHEELSED-- ---------- --------QK EVQQALAFIE ---------- ----------
[Human]HCoV-OC43    PCVASEQEES SEVLED---- --TLDDGPSV ETSDSQVE-- ---------- -----EDVEM SDFVDLESVI ---------- ----------
[Human]HCoV-NL63    FEEVEHVLSI KQPFSF---- --SFRDELGV RVLDQSDN-- ---------- ---------- ---------- ---------- ----------
```

```
                   271
[Feline]FCoV       EQEAEQPKPS LTP------- ---------- ---------- FKTTNLNGKI ILKQQDNNCW INACCYQLQA FDFFNHDLWD ---------G
[Porcine]TGEV      ----EQFNPS LPP------- ---------- ---------- FKTTNLNGKI ILKQGDNNCW INACCYQLQA FDFFNNEAWE ---------K
[Human]HCoV-HKU1   GDDVDDIESI YDFDTYKALL VFNDVYNDAL FVSYGSSVET ETYFKVNGLW SPTITHTNCW LRSVLLVMQK LPFKFKDLAI ENMWLSYKVG
[Bat]BtCoV/HKU4    ---------- ---------- ---------- ---------- ---------- ---------- ----AEVVET AEAQEPSVE- ---------S
[Human]SARS-CoV    ---------- ---------- ---------- ---------- ---------- ---------- ---------- ---------- ----------
[Human]SARS-CoV-2  ---------- ---------- ---------- ---------- ---------- ---------- ---------- ---------- ----------
[Human]MERS-CoV    ---------- ---------- ---------- ---------- ---------- ---------- ---------- ---------- ----------
[Bat]BtCoV/HKU9    ---------- ---------- ---------- ---------- ---------- ---------- ---------- ---------- ----------
[Human]HCoV-229E   ---------- ---------- ---------- ---------- ---------- -------NCW VNSVMLQIQL TGILDGDYAM Q--------F
[Avian]IBV         -KDALDVV-- ---------- ---------- ---------- ---------- ---------- ---------- ---------- ---------N
[Murine]MHV-A59    -LDELQTELN APADKTYEDV LAFDAVCSEA LSAFYAVPSD ETHFKVCGFY SPAIERTNCW LRSTLIVMQS LPLEFKDLEM QKLWLSYKAG
[Bat]BtCoV/HKU3    ---------- ---------- ---------- ---------- ---------- ---------- ---------- ---------- ----------
[Bat]BtCoV/512/2005 -DKPVVVKPD VFA------- ---------- ---------- FSYASYGGLK VLNQSSNNCW VSSALVQLQL TGLLDSDEMQ ---------L
[Human]HCoV-OC43   -QDYENVCFE FYT------- ---------- --------TE PEFVKVLGLY VPKATRNNCW LRSVLAVMQK LPCQFKDKNL QDLWVLYKQQ
[Human]HCoV-NL63   ---------- ---------- ---------- ---------- ---------- -------NCW ISTTLVQLQL TKLLDDSIEM Q--------L

                   361
[Feline]FCoV       FKKDDVMPFV DFCYAALTLK QGDSGDAEYL LETMLNDYST AKVTLSAKCG CGVKEIVLER TVFKLTPLRN EFKYGVCGDC KQINMCKFAS
[Porcine]TGEV      FKKGDVMDFV NLCYAATTLA RGHSGDAEYL LELMLNDYST AKIVLAAKCG CGEKEIVLER AVFKLTPLKE SFNYGVCGDC MQVNTCRFLS
[Human]HCoV-HKU1   YNQSFVDYLL TTIPKAIVLP QGGYVADFAY WFLNQFDINA YANWCCLKCG FSFDLNGLDA VFFYGDIVSH VCKCGHNMTL IAADL-----
[Bat]BtCoV/HKU4    IDSTPSTSTV VGENDLSVKP MSRVAETDDV LELETAVVGG ---------- ---------- ---------- ---------- ----------
[Human]SARS-CoV    ---------- ---------- ------IEP- ---------- ---------- ---------- ---------- ---------- ----------
[Human]SARS-CoV-2  -------GQQ DGSEDNQTTT IQTIVEVQPQ LEMEL----- ---------- ---------- ---------- ---------- ----------
[Human]MERS-CoV    VPVEDIAQVV IADTLQETPV VSDTVEVPPQ VVKL------ ---------- ---------- ---------- ---------- ----------
[Bat]BtCoV/HKU9    ------DNTI QDVDVATSMT TP-------- ---------- ---------- ---------- ---------- ---------- ----------
[Human]HCoV-229E   FKMGRVAKMI ERCYTAEQCI RGAMGDVGLC MYRLLKDLHT GFMVMDYKCS CTSGRLEESG AVLFCTPTKK AFPYGTCLNC NAPRMCTIRQ
[Avian]IBV         LPSGEETFVV NNCFEGAVKP LPQKVVD--- ---VLGDWGE AVDAQEQLCQ QEPLQHTFE- -----EPVEN STGSSKTMTE QVV-------
[Murine]MHV-A59    YDQCFVDKLV KSVPKSIILP QGGYVADFAY FFLSQCSFKA YANWRCLECD MELKLQGLDA MFFYGDVVSH MCKCGNSMTL LSADI-----
[Bat]BtCoV/HKU3    ---------- ---------- ---------- ---------- ---------- ---------- ---------- ---------- ----------
[Bat]BtCoV/512/2005 FNAGRVSPMV KRCYESQRAI FGSLGDVSAC LESLLKDRDG MSITCTIDCG CGPGVRVYEN AIFRFTPLKT AFPMGRCLIC SKTLMHTITQ
[Human]HCoV-OC43   YSQLFVDTLV NKIPANIVLP QGGYVADFAY WFLTLCDWQC VAYWKCIKCD LALKLKGLDA MFFYGDVVSH ICKCGESMVL IDVDV-----
[Human]HCoV-NL63   FKVGKVDSIV QKCYELSHLI SGSLGDSGKL LSELLKEKYT CSITFEMSCD CGKKFDDQVG CLFWIMPYTK LFQKGECCIC HKMQTYKLVS

                   451
[Feline]FCoV       VEGSGVFVHD RIEKQTPVSQ FIVTPTMHAV YTGTTQSGHY MIEDCIHDYC VDGMGI---K PRKHKFYTST LFLNANVMT- -----AKSKT
[Porcine]TGEV      VEGSGVFVHD ILSKQTPEAM FVVKPVMHAV YTGTTQNGHY MVDDIEHGYC VDGMGI---K PLKKRCYTST LFINANVMTR AEKPKQEFKV
[Human]HCoV-HKU1   ---------- ---PCTLHFS LFDDNFCAFC TPKKIFIAAC AVDV------ ---------- ---NVCHSVA VIGDEQIDGK FV---TKFSG
[Bat]BtCoV/HKU4    ---------- ---PVSDVTA IVTNDIVSVE QAQQCGVSSL PIQD------ ---------E ASENQVHQVS DLQGNELLC- -----SETKV
[Human]SARS-CoV    ---------- ---EPEPTPE EPVNQFTGYL K--------- ---------- ---------- ---------- ---------- ----------
[Human]SARS-CoV-2  ---------- ----TPVVQT IEVNSFSGYL K--------- ---------- ---------- ---------- ---------- ----------
[Human]MERS-CoV    ---------- ---PSEPQTI QPEVKEVAPV YE-------- ---------- ---------- ---------- ---------- -----ADTEQ
[Bat]BtCoV/HKU9    ---------- ---------- ------CGYT K--------- ---------- ---------- ---------- ---------- ----------
[Human]HCoV-229E   LQGTIIFVQQ KPEPVNPV-S FVVKPVCSSI FRGAVSCGHY QTNIYSQNLC VDGFGVNKIQ PWTNDALNTI CIKDADYN-- -----AKVEI
[Avian]IBV         ---------- -VEDQELPVV EQDQDVVVYT PTDLEVAKET AEEV------ ---------- ---DEFILIF AVPKEEVVSQ KDGAQIKQEP
[Murine]MHV-A59    ---------- ---PYTLHFG VRDDKFCAFY TPRKVFRAAC AVDV------ ---------- ---NDCHSMA VVEGKQIDGK VV---TKFIG
[Bat]BtCoV/HKU3    ---------- ---EPEPLPE EPVNQFVGYL K--------- ---------- ---------- ---------- ---------- ----------
[Bat]BtCoV/512/2005 MKGTGIFCRD A--TALDVDT LVVKPLCAAV YVGAQDGGHY LTNMYDANMA VDGHGR---- ---------- ---------- -----HPIKF
[Human]HCoV-OC43   ---------- ---PFTAHFA LKDKLFCAFI TKRIVYKAAC VVDV------ ---------- ---NDSHSMA VVDGKQIDDH RI---TSITS
[Human]HCoV-NL63   MKGTGVFVQD P--APIDIDA FPVKPICSSV YLGVKGSGHY QTNLYSFNKA IDGFGV---- ---------- ---------- -----FDIKN
```

```
                         541
[Feline]FCoV             MVEPPVPVED KCVE------ -DCQSPKDLI LPFYKAGKVS FYQGDLDVLI NFLEPDV--- -LVNAANGDL RHVGGVARAI DVFTGGKLTK
[Porcine]TGEV            EKVEQQPIVE ENKSSIEKEE IQSPKNDDLI LPFYKAGKLS FYQGALDVLI NFLEPDV--- -IVNAANGDL KHMGGVARAI DVFTGGKLTE
[Human]HCoV-HKU1         DKFDFIVGYG MSFS------ -MSSFELAQL YGLCITPNVC FVKGDIINVA RLVKADV--- -IVNPANGHM LHGGGVAKAI AVAAGKKFSK
[Bat]BtCoV/HKU4          EIVQPRQDLK PRRSR----- -KSKVDLSKY KHTVINNSVT LVLGDAIQIA SLLPKCI--- -LVNAANRHL KHGGGIAGVI NKASGGDVQE
[Human]SARS-CoV          ---------- ---------- ---------- -----LTDNVA IKCVDIVKEA QSANPMV--- -IVNAANIHL KHGGGVAGAL NKATNGAMQK
[Human]SARS-CoV-2        ---------- ---------- ---------- -----LTDNVY IKNADIVEEA KKVKPTV--- -VVNAANVYL KHGGGVAGAL NKATNNAMQV
[Human]MERS-CoV          TQSVTVKPKR LRKK------ -RNVDPLSNF EHKVITECVT IVLGDAIQVA KCYGESV--- -LVNAANTHL KHGGGIAGAI NAASKGAVQK
[Bat]BtCoV/HKU9          ---------- ---------- ---------- -----IAEHVY IKCADIVQEA RNYSYAV--- -LVNAANVNL HHGGGVAGAL NRATNNAMQK
[Human]HCoV-229E         SVTPIKNTVD TTPK------ -EEFVVKEKL NAFLVHDNVA FYQGDVDTVV NGVDFDF--- -IVNAANENL AHGGGLAKAL DVYTKGKLQR
[Avian]IBV               IQVVKPQREK KAKK------ -FKVKPATCE KPKFLEYKTC V--GDLTVVI AKALDEFKEF CIVNAANEHM THGSGVAKAI ADFCGLDFVE
[Murine]MHV-A59          DKFDFMVGYG MTFS------ -MSPFELAQL YGSCITPNVC FVKGDVIKVV RLVNAEV--- -IVNPANGRM AHGAGVAGAI AEKAGSAFIK
[Bat]BtCoV/HKU3          ---------- ---------- ---------- -----LTDNVA IKCIDIVKEA QSAKPTV--- -IVNAANTHL KHGGGVAGAL NKATNGAMQN
[Bat]BtCoV/512/2005      NTINTLCYKD VDWE------ -VSNGSCD-V KPFLTYKNIE FYQGELSALL S-VNHDF--- -VVNAANEQL SHGGGIAKAL DDLTKGELQV
[Human]HCoV-OC43         DKFDFIIGHG MSFS------ -MTTFEIAQL YGSCITPNVC FVKGDIIKVS KLVKAEV--- -VVNPANGHM VHGGGVAKAI AVAAGQQFVK
[Human]HCoV-NL63         SSVNTVCFVD VDFH------ -SVEIEAGEV KPFAVYKNVK FYLGDISHLV NCVSFDF--- -VVNAANENL LHGGGVARAI DILTEGQLQS

                         631
[Feline]FCoV             RSKEYLKSSK AIAPGNAVLF ENVLEHLS-V LNAVGPRNGD --SRVEGKLC NVYKAIAKCD GKI--LTPLI SVGIFKVKLE VSLQCLLKTV
[Porcine]TGEV            RSKDYLKKNK SIAPGNAVFF ENVIEHLS-V LNAVGPRNGD --SRVEAKLC NVYKAIAKCE GKI--LTPLI SVGIFNVRLE TSLQCLLKTV
[Human]HCoV-HKU1         ETAAMVKSKG VCQVGDCYVS TGGKLCKT-I LNIVGPDARQ DGRQSYVLLA RAYKHLNNYD -CC--LSTLI SAGIFSVPAD VSLTYLLGVV
[Bat]BtCoV/HKU4          ESDEYISNNG PLHVGDSVLL KGHGLADA-I LHVVGPDARN --NEDAALLK RCYKAFNKHT -IV--VTPLI SAGIFSVDPK VSFEYLLANV
[Human]SARS-CoV          ESDDYIKLNG PLTVGGSCLL SGHNLAKK-C LHVVGPNLNA --GEDIQLLK AAYENFNSQD -IL--LAPLL SAGIFGAKPL QSLQVCVQTV
[Human]SARS-CoV-2        ESDDYIATNG PLKVGGSCVL SGHNLAKH-C LHVVGPNVNK --GEDIQLLK SAYENFNQHE -VL--LAPLL SAGIFGADPI HSLRVCVDTV
[Human]MERS-CoV          ESDEYILAKG PLQVGDSVLL QGHSLAKN-I LHVVGPDARA --KQDVSLLS KCYKAMNAYP -LV--VTPLV SAGIFGVKPA VSFDYLIREA
[Bat]BtCoV/HKU9          ESSEYIKANG SLQPGGHVLL SSHGLASHGI LHVVGPDKRL --GQDLALLD AVYAAYTGFD -SV--LTPLV SAGIFGFTVE ESLCSLVKNV
[Human]HCoV-229E         LSKEHIGLAG KVKVGTGVMV ECDSLR---I FNVVGPRKG- --KHERDLLI KAYNTINNEQ GTP--LTPIL SCGIFGIKLE TSLEVLLDVC
[Avian]IBV               YCEDYVKKHG PQQRLVTPSF VKGIQC---V NNVVGPRHGD --NNLHEKLV AAYKNVL-VD GVVNYVVPVL SLGIFGVDFK MSIDAMREAF
[Murine]MHV-A59          ETSDMVKAQG VCQVGECYES AGGKLCKK-V LNIVGPDARG HGKQCYSLLE RAYQHINKCD -NV--VTTLI SAGIFSVPTD VSLTYLLGVV
[Bat]BtCoV/HKU3          ESDEYIRQNG PLTVGGSCLL SGHNLAEK-C LHVVGPNLNA --GEDVQLLK RAYENFNSQD -VL--LAPLL SAGIFGAKPL QSLKMCVEIV
[Bat]BtCoV/512/2005      LSNQYVSRNG SIKVGSGVLI KCKEHS---I LNVVGPRKG- --KHAAELLT KAYTFVFKQK GVP--LMPLL SVGIFKVPIT ESLAAFLACV
[Human]HCoV-OC43         ETTNMVKSKG VCATGDCYVS TGGKLCKT-V LNVVGPDART QGKQSYVLLE RVYKHFNNYD -CV--VTTLI SAGIFSVPSD VSLTYLLGTA
[Human]HCoV-NL63         LSKDYISSNG PLKVGAGVML ECEKFN---V FNVVGPRTG- --KHEHSLLV EAYNSILFEN GIP--LMPLL SCGIFGVRIE NSLKALFSCD

                         721
[Feline]FCoV             TD-------- ---------- ---------- ---------- ---------- ---------- ---------- ---------- ----------
[Porcine]TGEV            ND-------- ---------- ---------- ---------- ---------- ---------- ---------- ---------- ----------
[Human]HCoV-HKU1         DKQVILVSNN KEDFDII--- ---------- ---------- ---------- ---------- ---------- ---------- ----------
[Bat]BtCoV/HKU4          TTTTYVVVNN EDIYNTL--- ---------- ---------- ---------- ----ATPSKP ---------- ---------- ----------
[Human]SARS-CoV          RTQVYIAVND KALYEQVVMD YLD-----NL KPRV-EAPKQ EEPPNTEDSK TEEKSVVQKP VDVKPKIKAC IDEVTTTLEE TKFLTNKLLL
[Human]SARS-CoV-2        RTNVYLAVFD KNLYDKLVSS FLEMKSEKQV EQKIAEIPKE EVKPFITESK ----PSVEQR KQDDKKIKAC VEEVTTTLEE TKFLTENLLL
[Human]MERS-CoV          KTRVLVVVNS QDVYKSL--- ---------- ---------- ---------- ----TIVDIP ---------- ---------- ----------
[Bat]BtCoV/HKU9          ACTTYVVVYD RQLYERALAT SFDVPGPQS- ---------- ---------- ----SVQHVP AIDWAEAVEV QESIVDQVET PSL-------
[Human]HCoV-229E         NT-------- ---------- ---------- ---------- ---------- ---------- ---------- ---------- ----------
[Avian]IBV               ---------- ---------- ---------- ---------- ---------- ---------- ---------- ---------- ----------
[Murine]MHV-A59          TKNVILVSNN QDDFDVI--- ---------- ---------- ---------- ---------- ---------- ---------- ----------
[Bat]BtCoV/HKU3          RTQVYLAVND KSLYDQIVLD YLD-----SL KPKV-ESPNK EEEPKLEEPK AVQ-PVAEKP VDVKPKIKAC IDEVTTTLEE TKFLTNKLLL
[Bat]BtCoV/512/2005      GD-------- ---------- ---------- ---------- ---------- ---------- ---------- ---------- ----------
[Human]HCoV-OC43         KKQVVLVSNN QEDFDLI--- ---------- ---------- ---------- ---------- ---------- ---------- ----------
[Human]HCoV-NL63         IN-------- ---------- ---------- ---------- ---------- ---------- ---------- ---------- ----------
```

```
                 811
[Feline]FCoV     ---------- ---------- ---------- ---------- ---------- ---------- ---------- ------RDLN VFVYTDQERV
[Porcine]TGEV    ---------- ---------- ---------- ---------- ---------- ---------- ---------- ------RGLN VFVYTDQERQ
[Human]HCoV-HKU1 ---------- ---------- ---------- ---------- ---------- ---------- ---------- ------QKCQ ITSVVGTKAL
[Bat]BtCoV/HKU4  ---------- ---------- ---------- ---------- ---------- ---------- ---------- ------DGLV -YSFEGWRG-
[Human]SARS-CoV  FADINGKLYH DSQNMLRGED MSFLEKDAPY MVGDVITSGD ITCVVIPSKK AGGTTEMLSR ALKKVPVDEY ITTYPGQGCA GYTLEEAKT-
[Human]SARS-CoV-2 YIDINGNLHP DSATLVSDID ITFLKKDAPY IVGDVVQEGV LTAVVIPTKK AGGTTEMLAK ALRKVPTDNY ITTYPGQGLN GYTVEEAKT-
[Human]MERS-CoV  ---------- ---------- ---------- ---------- ---------- ---------- ---------- ------QSLT -FSYDGLRG-
[Bat]BtCoV/HKU9  ---------- --------GA VDTVDSNADS GLNETARSPE NVVGSVPDDV VADVESCVRD LVRQVVKKVK RDKRPPPIVP QQTVEQQPQ-
[Human]HCoV-229E ---------- ---------- ---------- ---------- ---------- ---------- ---------- ------KEVK VFVYTDTEVC
[Avian]IBV       ---------- ---------- ---------- ---------- ---------- ---------- ---------- ------EGCT IRVL------
[Murine]MHV-A59  ---------- ---------- ---------- ---------- ---------- ---------- ---------- ------EKCQ VTSVAGTKAL
[Bat]BtCoV/HKU3  FADINGKLYQ DSQNMLRGED MSFLEKDAPY IVGDVITSGD ITCVIIPAKK SGGTTEMLAR ALKEVPVAEY ITTYPGQGCA GYTLEEAKT-
[Bat]BtCoV/512/2005 ---------- ---------- ---------- ---------- ---------- ---------- ---------- ------RVCK CFCYTDKERL
[Human]HCoV-OC43 ---------- ---------- ---------- ---------- ---------- ---------- ---------- ------SKCQ ITAVEGTKKL
[Human]HCoV-NL63 ---------- ---------- ---------- ---------- ---------- ---------- ---------- ------KPLQ VFVYSSNEEQ

                 901
[Feline]FCoV     TIENFFN-G- ---------- ---------- ---------- ---------- ---------- ---------- ---------- ----------
[Porcine]TGEV    TIENFFS-C- ---------- ---------- ---------- ---------- ---------- ---------- ---------- ----------
[Human]HCoV-HKU1 AVRLTAN--- ---------- ---------- ---------- ---------- ---------- ---------- ------VGRV IKFETDAYKL
[Bat]BtCoV/HKU4  TVRTAKNYGF TCF------- ---------- ---------- -------ICT EYSANVKFL- -RTKGVDTTK KIQTVDGVSY YLYSARDALT
[Human]SARS-CoV  ALKKCKS-AF YVLPSEAPNA KEEILGTVSW NLREMLAHAE ETRKLMPICM DVRAIMATIQ RKYKGIKIQE GIVDY-GVRF FFYTSKEPVA
[Human]SARS-CoV-2 VLKKCKS-AF YILPSIISNE KQEILGTVSW NLREMLAHAE ETRKLMPVCV ETKAIVSTIQ RKYKGIKIQE GVVDY-GARF YFYTSKTTVA
[Human]MERS-CoV  AIRKAKDYGF TVF------- ---------- ---------- -------VCT DNSANTKVL- -RNKGVDYTK KFLTVDGVQY YCYTSKDTLD
[Bat]BtCoV/HKU9  EISSPGD-CN TVL------- ----VDVVSM SFSAMVNFGK EKGLLIPVVI DYPAFLKVL- ---KRFSPKE GLFSSNGYEF YGYSRDKPLH
[Human]HCoV-229E KVKDFVS-GL VNV------- ---------- ---------- ---QKVEQPK IEPKPVSVIK VAPKPYRV-- ------DGKF SYFTEDLLCV
[Avian]IBV       LFSLSQE--- ---------- ---------- ---------- ---------- ---------- ---------- ------HIDY FDVTCK----
[Murine]MHV-A59  SLQLAKN--- ---------- ---------- ---------- -------LCR DVKFV----- ---------- ---------- -----TNACS
[Bat]BtCoV/HKU3  ALKKCKS-AF YVLPSETPNE KEEVLGTVSW NLREMLAHAE ETRKLMPICL DVRAIMATIQ RKYKGIKVQE GIVDY-GVRF FFYTSKEPVA
[Bat]BtCoV/512/2005 AIQNFVT-SF QTE------- ---------- ---QPVEPLP VIQEVKGVQL EKPVPDVKVE NPCEPFRIEG ------DAKF YDLTPSMVQS
[Human]HCoV-OC43 AARLSFN--- ---------- ---------- ---------- ---------- ---------- ---------- ------VGRS IVYETDANKL
[Human]HCoV-NL63 AVLKFLD-GL DLT------- ---------- ---------- ------PVID DVDVV----- ---KPFRV-- ------EGNF SFFDCGVNAL

                 991
[Feline]FCoV     ---------- ---------- ---------- ---------- ---------- ---------- ---------- ---------- ----------
[Porcine]TGEV    ---------- ---------- ---------- ---------- ---------- ---------- ---------- ---------- ----------
[Human]HCoV-HKU1 FLSGDDCFVS NSSVIQEVLL LRHDIQLNND VRDYLLSKMT SLP------- ---------- ---------- ---------- --KDWRLINK
[Bat]BtCoV/HKU4  DVIAAANG-C SGICAMPFGY VTHGLDLAQS GNYVRQVKVP YVCLLASKEQ IPIMNS--D- --VAIQTPET AFINNVTSNG GYHSWHLVSG
[Human]SARS-CoV  SIITKLNSLN EPLVTMPIGY VTHGFNLEEA ARCMRSLKAP AVVSVSSPDA VTTYNGYLT- --SSSKTSEE HFVETVSLAG SYRDWSY-SG
[Human]SARS-CoV-2 SLINTLNDLN ETLVTMPLGY VTHGLNLEEA ARYMRSLKVP ATVSVSSPDA VTAYNGYLT- --SSSKTPEE HFIETISLAG SYKDWSY-SG
[Human]MERS-CoV  DILQQANK-S VGIISMPLGY VSHGLDLIQA GSVVRRVNVP YVCLLANKEQ EAILMS--E- --DVKLNPSE DFIKHVRTNG GYNSWHLVEG
[Bat]BtCoV/HKU9  EVSKDLNSLG RPLIMIPFGF IVNGQTLAVS AVSMRGLTVP HTVVVPSESS VPLYRAYFNG VFSGDTTAVQ DFVVDILLNG A-RDWDVLQT
[Human]HCoV-229E ADDKPIVLFT DSMLTLDDRG LALDNALSGV LSAAIKDCVD INKAIPSGNL IKFDIGSVV- ---------- VYMCVVPSEK D-------KH
[Avian]IBV       ---------- ---------- ---------- ---------- ---------- ---------- ---------- ---------- ----------
[Murine]MHV-A59  SLFSESCFVS SYDVLQEVEA LRHDIQLDDD ARVFVQANMD CLP------- ---------- ---------- ---------- --TDWRLVNK
[Bat]BtCoV/HKU3  SIITKLNSLN EPLVTMPIGY VTHGLNLEEA ARCMRSLKAP AVVSVSSPDA VTAYNGYLT- --SSSKTPEE YFVETTSLAG SYRDWSY-SG
[Bat]BtCoV/512/2005 LQVTRLVSFT NSDLCLGSFV RDCDGYVQGS LGGAIANYKK SNPVLPAGNC VTLKCDGFI- ---------S FTFVILPKEG D-------TN
[Human]HCoV-OC43 ILINDVAFVS TFNVLQDVLS LRHDIALDDD ARTFVQSNVD VLP------- ---------- ---------- ---------- --EGWRVVNK
[Human]HCoV-NL63 DGDIY-LLFT NSILMLDKQG QLLDTKLNGI LQQAALDYLA TVKTVPAGNL VKLFVESCT- ---------- IYMCVVPSIN D-------LS
```

```
                    1081
[Feline]FCoV        ---------- ---------- ---------- ---------- ---------- ---------- TIPIKVTEDT VNQKRVSVAL DKTYGEQLKG
[Porcine]TGEV       ---------- ---------- ---------- ---------- ---------- ---------- SIPVNVTEDN VNHERVSVSF DKTYGEQLKG
[Human]HCoV-HKU1    FDVINGVKTV KYFECPNSIY ICSQG-KDFG YVCDGSFYK- -ATVNQVCVL LAKK------ -IDVLLTVDG VNFKSISLTV GEVFGKIL-G
[Bat]BtCoV/HKU4     DLIVKDVCYK KLLH-WSGQT ICYAD-NKFY VVKNDVALPF -SDLEACRAY LTSR-AAQQV NIEVLVTIDG VNFRTVILND TTTFRKQL-G
[Human]SARS-CoV     QRTELGV--- EFLK-RGDKI VYHTLESPVE FHLDGEVLSL -DKL---KSL LSLR-EVK-- TIKVFTTVDN TNLHTQLVDM SMTYGQQF-G
[Human]SARS-CoV-2   QSTQLGI--- EFLK-RGDKS VYYTS-NPTT FHLDGEVITF -DNL---KTL LSLR-EVR-- TIKVFTTVDN INLHTQVVDM SMTYGQQF-G
[Human]MERS-CoV     ELLVQDLRLN KLLH-WSDQT ICYKD-SVFY VVKNSTAFPF -ETLSACRAY LDSR-TTQQL TIEVLVTVDG VNFRTVVLNN KNTYRSQL-G
[Bat]BtCoV/HKU9     TCTVDRKVYK TICK-RGNTY LCFDD-TNLY AITGDVVLKF -ATVSKARAY LETKLCAPEP LIKVLTTVDG INYSTVLVST AQSYRAQI-G
[Human]HCoV-229E    LDNNVQRCTR KLNRLMCDIV CTIPADYILP LVLSSLTCNV -SFVGELKAA EAKV------ -ITIKVTEDG VNVHDVTVTT DKSFEQQV-G
[Avian]IBV          ---------- ---------- ---------- ---------- ---------- ---------- QKTIYLTEDG VKYRSIVLKP GDSLG-QF-G
[Murine]MHV-A59     FDSVDGVRTI KYFECPGGIF VSSQG-KKFG YVQNGSFKE- -ASVSQIRAL LANK------ -VDVLCTVDG VNFRSCCVAE GEVFGKTL-G
[Bat]BtCoV/HKU3     QRTELGV--- EFLK-RGDKI VYHTTGSPIE FHLDGEVLPL -DKL---KSL LSLR-EVK-- TIKVFTTVDN TNLHTHIVDM SMTYGQQF-G
[Bat]BtCoV/512/2005 YEKNFNRAIA KFLKLKGSLL VVVEDSSVFN KISHASVAGY VAKPALVDTL FEAK------ PVQVVVTQDQ RSFHTVELST SQTYGQQL-G
[Human]HCoV-OC43    FYQINGVRTV KYFECTGGID ICSQD-KVFG YVQQGIFNK- -ATVAQIKAL FLDK------ -VDILLTVDG VNFTNRFVPV GESFGKSL-G
[Human]HCoV-NL63    FDKNLGRCVR KLNRLKTCVI ANVPAIDVLK KLLSSLTLTV -KFVVESNVM DVND-CFKND NVVLKITEDG INVKDVVVES SKSLGKQL-G

                    1171
[Feline]FCoV        TVVIKDKDVT NQLPSVSDVG EKVVK---AL DVDWN----- AYYGFPNAA- --AFSASSHD AYEFDVVTHN NFIVHKQTDN NCWVNAICLA
[Porcine]TGEV       TVVIKDKDVT NQLPSAFDVG QKVIK---AI DIDWQ----- AHYGFRDAA- --AFSASSHD AYKFEVVTHS NFIVHKQTDN NCWINAICLA
[Human]HCoV-HKU1    NVFCDGIDVT KLKCSDFYAD KILYQYENLS LADISAVQ-- SSFGFDQQQL L-AYYNFLTV CK-WSVVVNG PFFSFEQSHN NCYVNVACLM
[Bat]BtCoV/HKU4     ATFYKGVDIS DAFPTVKMGG ESLFVADNLS ESEKVVLK-- EYYGTSDVTF LQRYYSLQPL VQQWKFVVHD GVKSLKLSNY NCYINATIMM
[Human]SARS-CoV     PTYLDGADVT KIKPHVNHEG KTFFV---LP SDDTLRSEAF EYYHTLDESF LGRYMSALNH TKKWKFPQVG GLTSIKWADN NCYLSSVLLA
[Human]SARS-CoV-2   PTYLDGADVT KIKPHNSHEG KTFYV---LP NDDTLRVEAF EYYHTTDPSF LGRYMSALNH TKKWKYPQVN GLTSIKWADN NCYLATALLT
[Human]MERS-CoV     CVFFNGADIS DTIPDEKQNG HSLYLADNLT ADETKALK-- ELYGPVDPTF LHRFYSLKAA VHKWKMVVCD KVRSLKLSDN NCYLNAVIMT
[Bat]BtCoV/HKU9     TVFCDGHDWS NKNPMPTDEG THLYKQDNFS SAEVTAIR-- EYYGVDDSNI IARAMSIRKT VQTWPYTVVD GRVLLAQRDS NCYLNVAISL
[Human]HCoV-229E    VIADKDKDLS GAVPSDLNTS ELLTK---AI DVDWV----- EFYGFKDAV- --TFATVDHS AFAYESAVVN GIRVLKTSDN NCWVNAVCIA
[Avian]IBV          QVYAKNKIV- -FTADDVEDK EILYV----P TTDKSIL--- EYYGLDAQKY V---IYLQTL AQKWNVQYRD NFLILEWRDG NCWISSAIVL
[Murine]MHV-A59     SVFCDGINVT KVRCSAIYKG KVFFQYSDLS EADLVAVK-- DAFGFDEPQL L-KYYTMLGM CK-WPVVVCG NYFAFKQSNN NCYINVACLM
[Bat]BtCoV/HKU3     PTYLDGADVT KIKPHVNHEG KTFFV---LP SDDTLRSEAF EYYHTIDESF LGRYMSALNH TKKWKFPQVG GLTSIKWADN NCYLSSVLLA
[Bat]BtCoV/512/2005 DCVVEDKKVT NLKP--VSKD KVVSV---VP NVDWD----- KHYGFVDAG- --IFHTLDHT MFVFDNNVVN GKRVLRTSDN NCWINAVCLQ
[Human]HCoV-OC43    NVFCDGVNVT KHKCDINYKG KVFFQFDNLS SEDLKAVR-- SSFNFDQKEL L-AYYNMLVN CFKWQVVVNG KYFTFKQANN NCFVNVSCLM
[Human]HCoV-NL63    VVSDGVDSFE GVLP--INTD TVLSV---AP EVDWV----- AFYGFEKAA- --LFASLDVK PYGYPNDFVG GFRVLGTTDN NCWVNATCII

                    1261
[Feline]FCoV        LQRL-KPTWK FPGVKSLWDA FLTRKTAGFV HMLYHISGLT KGQPGDAELT LHKLVDLMSS D-SAVTVTHT TACDKC---- -AKVETFTGP
[Porcine]TGEV       LQRL-KPQWK FPGVRGLWNE FLERKTQGFV HMLYHISGVK KGEPGDAELM LHKLGDLMDN D-CEIIVTHT TACDKC---- -AKVEKFVGP
[Human]HCoV-HKU1    LQHI-NLKFN KWQWQEAWYE FRAGRPHRLV ALVLAKGHFK FDEPSDATDF IRVVLKQADL S--GAICELE LICD-CGIK- QESRVGVDAV
[Bat]BtCoV/HKU4     IDMLHDIKFV VPALQNAYLR YKGGDPYDFL ALIMAYGDCT FDNPDDEAKL LHTLLAKAEL T-VSAKMVWR EWCTVCGIR- DIEYTGMRAC
[Human]SARS-CoV     LQQL-EVKFN APALQEAYYR ARAGDAANFC ALILAYSNKT VGELGDVRET MTHLLQHANL E--SAKRVLN VVCKHCGQK- TTTLTGVEAV
[Human]SARS-CoV-2   LQQI-ELKFN PPALQDAYYR ARAGEAANFC ALILAYCNKT VGELGDVRET MSYLFQHANL D--SCKRVLN VVCKTCGQQ- QTTLKGVEAV
[Human]MERS-CoV     LDLLKDIKFV IPALQHAFMK HKGGDSTDFI ALIMAYGNCT FGSPDDASRL LHTVLAKAEL C-CSARMVWR EWCNVCGIK- DVVLQGLKAC
[Bat]BtCoV/HKU9     LQDI-DVSFS TPWVCRAYDA LKGGNPLPMA EVLIALGKAT PGVSDDAHMV LSAVLNHGTV T---ARRVMQ TVCEHCGVS- QMVFTGTDAC
[Human]HCoV-229E    LQYS-KPHFI SQGLDAAWNK FVLGDVEIFV AFVYYVARLM KGDKGDAEDT LTKLSKYLAN E-AQVQLEHY SSCVECDAKF KNSVASINSA
[Avian]IBV          LQAA-KIRFK GF-LTEAWAK LLGGDPTDFV AWCYASCTAK VGDFSDANWL LANLAEHFDA DYTNAFLKKR VSCN-CGIK- SYELRGLEAC
[Murine]MHV-A59     LQHL-SLKFP KWQWQEAWNE FRSGKPLRFV SLVLAKGSFK FNEPSDSIDF MRVVLREADL S--GATCNLE FVCK-CGVK- QEQRKGVDAV
[Bat]BtCoV/HKU3     LQQV-EVKFN APALQEAYYR ARAGDAANFC ALILAYSNKT VGELGDVRET MTHLLQHANL E--SAKRVLN VVCKHCGQK- TTTLKGVEAV
[Bat]BtCoV/512/2005 LQFA-NAKFK PKGLQQLWES YCTGDVAMFV HWLYWITGVE KGEPSDAENT LNIISRFLKP Q--GSVEMLR ATSTTCDGT- CSTKRVVSTP
[Human]HCoV-OC43    LQSL-HLTFK IVQWQEAWLE FRSGRPARFV ALVLAKGGFK FGDPADSRDF LRVVFSQVDL T--GAICDFE IACK-CGVK- QEQRTGLDAV
[Human]HCoV-NL63    LQYL-KPTFK SKGLNVLWNK FVTGDVGPFV SFIYFITMSS KGQKGDAEEA LSKLSEYLIS D-SIVTLEQY STCDIC---- KSTVVEVKSA
```

```
                         1351
[Feline]FCoV             VV-AAPLLVC GTDE-I---C VHGVHVNVKV TSIRGTVAIT SLI------- GPVVGDVIDA T-----GYIC YTG-LNSRGH YTYYDNRNGL
[Porcine]TGEV            VV-AAPLAIH GTDET----C VHGVSVNVKV TQIKGTVAIT SLI------- GPIIGEVLEA T-----GYIC YSG-SNRNGH YTYYDNRNGL
[Human]HCoV-HKU1         MH-FGTLAKT DLFNGYKIGC NCAG-RIVHC TKLNVPFLIC SN-------- TPLSKDLPDD V----VAANM FMG--VGVGH YTHLKCGSPY
[Bat]BtCoV/HKU4          VY-AGVNSME ELQSVFNETC VCGSVKHRQL VEHSAPWLLV S--------- GLNEVKVSTS TDPIYRAFNV FQGVETSVGH YVHIRVKDGL
[Human]SARS-CoV          MY-MGTLSYD NLKTGVSIPC VCGRDATQYL VQQESSFVMM SA-------- PPAEYKLQQG T---FLCANE YTG-NYQCGH YTHITAKETL
[Human]SARS-CoV-2        MY-MGTLSYE QFKKGVQIPC TCGKQATKYL VQQESPFVMM SA-------- PPAQYELKHG T---FTCASE YTG-NYQCGH YKHITSKETL
[Human]MERS-CoV          CY-VGVQTVE DLRARMTYVC QCGGERHRQI VEHTTPWLLL SG-------- TPNEKLVTTS TAPDFVAFNV FQGIETAVGH YVHARLKGGL
[Bat]BtCoV/HKU9          TF-YGSVVLD DLYAPVSVVC QCGRPAIRYV SEQKSPWLLM SC-------- TPTQVPLDTS G--IWKTAIV FRG-PVTAGH YMYA-VNGTL
[Human]HCoV-229E         IV-CASVKRD GVQVGY---C VHGIKYYSRV RSVRGRAIIV SVEQLE---- PCAQSRLLSG V-----AYTA FSG-PVDKGH YTVYDTAKKS
[Avian]IBV               IQPVRATNLL HFKTQYSNCP TCGANNTDEV IEASLPYLLL FATD------ GPATVDCDED A----VGTVV FVG-STNSGH C--YTQAAGQ
[Murine]MHV-A59          MH-FGTLDKG DLVRGYNIAC TCGS-KLVHC TQFNVPFLIC SN-------- TPEGRKLPDD V----VAANI FTG--GSVGH YTHVKCKPKY
[Bat]BtCoV/HKU3          MY-MGTLSYD ELKTGVSIPC VCGRNATQYL VQQESSFVMM SA-------- PPAEYKLQQG A---FLCANE YTG-NYQCGH YTHITAKETL
[Bat]BtCoV/512/2005      VV-NASVLKV GLDDGN---C VHGLPLVDRV VSVNGTVIIT NVGDTPGKPV VATENLLLDG V-----SYTV FQDSTTGVGH YTVFDKEAKL
[Human]HCoV-OC43         MH-FGTLSRE DLEIGYTVDC SCGK-KLIHC VRFDVPFLIC SN-------- TPASVKLPKG V----GSANI FIG--DNVGH YVHVKCEQSY
[Human]HCoV-NL63         IV-CASVLKD GCDVGF---C PHRHKLRSRV KFVNGRVVIT NVGEPI---- ISQPSKLLNG I-----AYTT FSG-SFDNGH YVVYDAANNA

                         1441
[Feline]FCoV             --MVDADKAY HFEKNLLQVT TAIA------ ---------- ---------- ---------- SNFVANTPKK EIMPKTQAKE SKAKESN---
[Porcine]TGEV            --VVDAEKAY HFNRDLLQVT TAIA------ ---------- ---------- ---------- SNFVVKKPQA EERPKNCAFN KVAASPKI--
[Human]HCoV-HKU1         -QHYDACSVK KYTGVSGCLT DCLYLKNLTQ TFTSMLTNYF LDDVEMVAYN PDLSQYYCDN GKYYTKPIIK AQFKPFAKVD GVYTNFKL-V
[Bat]BtCoV/HKU4          FYKYDSGSLT KTSDMKCKMT SVWY-PTVRY TADCNVVVYD LDGVTKVEVN PDLSNYYMKD GKYYTSKPTI KYSPATILPG SVYSNSCL-V
[Human]SARS-CoV          -YRIDGAHLT KMSEYKGPVT DVFY-KETSY TTTIKPVSYK LDGVTYTEIE PKLDGYYKKD NAYYTEQP-I DLVPTQPLPN ASFDNFKL-T
[Human]SARS-CoV-2        -YCIDGALLT KSSEYKGPIT DVFY-KENSY TTTIKPVTYK LDGVVCTEID PKLDNYYKKD NSYFTEQP-I DLVPNQPYPN ASFDNFKF-V
[Human]MERS-CoV          ILKFDSGTVS KTSDWKCKVT DVLF-PGQKY SSDCNVVRYS LDGNFRTEVD PDLSAFYVKD GKYFTSEPPV TYSPATILAG SVYTNSCL-V
[Bat]BtCoV/HKU9          ISVYDANTRR RTSDLKLPAT DILY-GPTSF TSDSKVETYY LDGVKRTTID PDFSKYVKRG DYYFTTAP-I EVVAAPKLV- TSYDGFYLSS
[Human]HCoV-229E         --MYDGDRFV KHDLSLLSVT SVVM------ ---------- ---------- ---------V GGYVA----- ---------- ----------
[Avian]IBV               --AFDNLAKD RKFGKKSPYI TAMY-TRFAF KNETSLPVAK QSKGKSKSVK EDVSNLATSS KASFDNLTDF EQW----YDS NIYESLKV--
[Murine]MHV-A59          -QLYDACNVN KVSEAKGNFT DCLYLKNLKQ TFSSVLTTFY LDDVKCVEYK PDLSQYYCES GKYYTKPIIK AQFRTFEKVD GVYTNFKL-V
[Bat]BtCoV/HKU3          -YRVDGAHLT KMSEYKGPVT DVFY-KETSY TTAIKPVSYK LDGVTYTEIE PKLDGYYKKG NAYYTEQP-I DLVPTQPMPN ASFDNFKL-T
[Bat]BtCoV/512/2005      --MFDGDVLK PCDLNVSPVT SVVV------ ---------- ---------- -------CNN KKIVVQDP-- ---------- ----------
[Human]HCoV-OC43         -QLYDASNVK KVTDVTGKLS DCLYLKNLKQ TFKSVLTTYY LDDVKKIEYK PDLSQYYCDG GKYYTQRIIK AQFKTFEKVD GVYTNFKL-I
[Human]HCoV-NL63         --VYDGARLF SSDLSTLAVT AIVV------ ---------- ---------- ---------V GGCVTSNV-- ---------- ----------

                         1531
[Feline]FCoV             ---------- ---------- ---------- ---------- ---------- ---------- ---------- ---------- -----TARVF
[Porcine]TGEV            ---------- ---------- ---------- ---------- ---------- ---------- ---------- ---------- ----------
[Human]HCoV-HKU1         GHDICAQ--- ----LNDKLG FNVDLPFV-E YKVTVWPVAT GDVVLASDDL YVKRYFKGCE TFGKPVIWFC HDEA------ -----SLNSL
[Bat]BtCoV/HKU4          GVDGTPGSDT ISKFFNDLLG FDETKPISKK LTYSLLPNED GDVLLSEFSN YNPVYKKGVM LKGKPILWVN NGVCDSALNK PNRA-SLRQL
[Human]SARS-CoV          CSNTKFADD- ----LNQMTG F--TKPASRE LSVTFFPDLN GDVVAIDYRH YSASFKKGAK LLHKPIVWHI NQATTKTTFK PNTW-CLRCL
[Human]SARS-CoV-2        CDNIKFADD- ----LNQLTG Y--KKPASRE LKVTFFPDLN GDVVAIDYKH YTPSFKKGAK LLHKPIVWHV NNATNKATYK PNTW-CIRCL
[Human]MERS-CoV          SSDGQPGGDA ISLSFNNLLG FDSSKPVTKK YTYSFLPKED GDVLLAEFDT YDPIYKNGAM YKGKPILWVN KASYDTNLNK FNRA-SLRQI
[Bat]BtCoV/HKU9          CQNPQLAES- ----FNKAIN ATKTGPMK-- -LLTMYPNVA GDVVAIS-DD NVVAHPYGSL HMGKPVLFVT ---------R PNTWKKLVPL
[Human]HCoV-229E         ---------- ---------- ---------- ---------- ---------- ---------- ---------- ---------- ----------
[Avian]IBV               ---------- ---------- ---------- ---------- ---------- ---------- ---------- ---------- ----------
[Murine]MHV-A59          GHS--IAEK- ----LNAKLG FDCNSPFV-E YKITEWPTAT GDVVLASDDL YVSRYSSGCI TFGKPVVWLG HEEA------ -----SLKSL
[Bat]BtCoV/HKU3          CSNTKFADD- ----LNQMTG F--KKPASRE LTVTFFPDLN GDVVAIDYRH YSTSFKKGAK LVHKPILWHI NQTTNKTTYK PNIW-CLRCL
[Bat]BtCoV/512/2005      ---------- ---------- ---------- ---------- ---------- ---------- ---------- ---------- ----------
[Human]HCoV-OC43         GHT--VCDS- ----LNSKLG FDSSKEFV-E YKITEWPTAT GDVVLANDDL YVKRYERGCI TFGKPVIWLS HEKA------ -----SLNSL
[Human]HCoV-NL63         ---------- ---------- ---------- ---------- ---------- ---------- ---------- ---------- ----------
```

```
                    1621
[Feline]FCoV        SEVEENPKNI VRKEKLLAIE SGV------- ---------- ---------- ---------- ---------- ---------- ----DYTITT
[Porcine]TGEV       ---------- VQEQKLLAIE SGA------- ---------- ---------- ---------- ---------- ---------- ----NYALTE
[Human]HCoV-HKU1    TYFNKPSFKS ENRYSVLSVD SVS------E ESQGNVVTSV MESQ------ ---------- ---------- ---------- ----ISTKEV
[Bat]BtCoV/HKU4     --YDVAPIVL DNKYTVLQDN TSQ-----LV EHNVPVVDDV PITT------ ---------- ---------- ---------- ----RKLIEV
[Human]SARS-CoV     --WSTKPVDT SNSFEVLAVE DTQGMDNLAC ESQQPTSEEV VENP------ ---------- ---------- ---------- ----TIQKEV
[Human]SARS-CoV-2   --WSTKPVET SNSFDVLKSE DAQGMDNLAC EDLKPVSEEV VENP------ ---------- ---------- ---------- ----TIQKDV
[Human]MERS-CoV     --FDVAPIEL ENKFTPLSVE STP------- --VEPPTVDV VALQ------ ---------- ---------- ---------- ----QEMTIV
[Bat]BtCoV/HKU9     --LSTVVVNT PNTYDVLAVD ---------- --PLPVNNET S--------- ---------- --------EE PISV------ ----K--API
[Human]HCoV-229E    ------PVNT VKP------- ---------- ---------- ---------- ---------- ---------- ---------- ----KPVINQ
[Avian]IBV          ---QESPDNF DKYVSFTTKE DSK------- ---LPLTLKV ---------- ---------- ---------- ---------- ----RGIKSV
[Murine]MHV-A59     TYFNRPSVVC ENKFNVLPVD VSE------P TDKGPVPAAV LVTGVPGADA SAGAGIAKEQ KACASASVED QVVTEVRQEP SVSAADVKEV
[Bat]BtCoV/HKU3     --WSTKPVDT SNSFEVLVVE DTQGMDNLAC ESQTTTSEEV VENP------ ---------- ---------- ---------- ----TVQKEI
[Bat]BtCoV/512/2005 --VKRVELDA SKFLDTMNVA S--------- ---------- ---------- ---------- ---------- ---------- ----------
[Human]HCoV-OC43    TYFNRPLLVD DNKFDVLKVD DVD------- -DSGDSSESG AKET------ ---------- ---------- ---------- ----KEINII
[Human]HCoV-NL63    ------PTIV SEKISVM--- ---------- ---------- ---------- ---------- ---------- ---------- --------DK

                    1711
[Feline]FCoV        LGKYADVFFM AGDKILRF-- ---------- ---------- ---------- ---------- ---------- ---------- ----------
[Porcine]TGEV       FGRYADMFFM AGDKILRL-- ---------- ---------- ---------- ---------- ---------- ---------- ----------
[Human]HCoV-HKU1    KLKGVRKTVK IEDAIIVNDE NSSIKVVKSL SLVDVWDMYL T-GCDYVVWV ANELSRLVKS PTVREY---- ------IRYG IKPITIPIDL
[Bat]BtCoV/HKU4     KCKGLNKPFV KGNFSFVNDP NG-VTVVDTL GLTELRALYV DINTRYIVLR DNNWSSLFKL HTVESGDLQI VAAGGSVTRR ARVLLGASSL
[Human]SARS-CoV     IECDVKTTEV VGNVILKPSD EG-VKVTQEL GHEDLMAAYV E-NTSITIKK PNELSLALGL KTIATH---G IAAINSVPWS KILAYVKPFL
[Human]SARS-CoV-2   LECNVKTTEV VGDIILKPAN NS-LKITEEV GHTDLMAAYV D-NSSLTIKK PNELSRVLGL KTLATH---G LAAVNSVPWD TIANYAKPFL
[Human]MERS-CoV     KCKGLNKPFV KDNVSFVADD SG-TPVVEYL SKEDLHTLYV DPKYQVIVLK DNVLSSMLRL HTVESGDINV VAASGSLTRK VKLLFRASFY
[Bat]BtCoV/HKU9     PLYGLKATMV LNGTTYVPGN KGHLLCLKEF TLTDLQTFYV EGVQPFVLLK ASHLSKVLGL RVSDSS---- -LHVNHLSKG VVYAYAATRL
[Human]HCoV-229E    LDEKAQKFFD FGDFLIHN-- ---------- ---------- ---------- ---------- ---------- ---------- ----------
[Avian]IBV          VDFRSKDGFI YKLTPDTDEN SKAPVYYPVL DAISLKAIWV EGNANFVVGH PNYYSKSLHI PTFWENAENF VKMGDKIGGV TMGLWRAEHL
[Murine]MHV-A59     KLNGVKKPVK VEGSVVVNDP TSETKVVKSL SIVDVYDMFL T-GCKYVVWT ANELSRLVNS PTVREY---- ------VKWG MGKIVTPAKL
[Bat]BtCoV/HKU3     IECDVKTTEV VGNVILKPSE EG-VKVTQEL GHEDLMAAYV E-ETSITIKK PNELSLALGL KTLATH---G AAAINSVPWS KILAYVKPFL
[Bat]BtCoV/512/2005 -----EKFFT FGDFVSRN-- ---------- ---------- ---------- ---------- ---------- ---------- ----------
[Human]HCoV-OC43    KLSGVKKPFK VEDSVIVNDD TSETKYVKSL SIVDVYDMWL T-GCKYVVRT ANALSRAVNV PTIRKF---- ------IKFG MTLVSIPIDL
[Human]HCoV-NL63    LDTGAQKFFQ FGDFVMNN-- ---------- ---------- ---------- ---------- ---------- ---------- ----------

                    1801
[Feline]FCoV        ---------- ---------- ---------- --------LL EVFKYLLVVF M-CL--RKSK MPKVKVKPPH VFRNLGAKVR TLNY------
[Porcine]TGEV       ---------- ---------- ---------- --------LL EVFKYLLVLF M-CL--RSTK MPKVKVKPPL AFKDFGAKVR TLNY------
[Human]HCoV-HKU1    ---LCLRDDN QTLLVPKIFK ARAIE----- --------FY GFLKWLFIYV FSLL--HFTN DK------TI FYTTEIASKF TFNLFCLA--
[Bat]BtCoV/HKU4     FASFAKITVT ATTAACKTAG RGFCK---FV VNYGVLQNMF VFLKMLFFLP FNYL--WPKK QPTVDIGVSG LRTAGIVTTN IVKQCGTAAY
[Human]SARS-CoV     GQAAITTSNC AKRLAQRV-- ---------- --------FN NYMPYVFTLL FQLC--TFTK STNSRIRASL PTT---IAKN SVKSVAKLCL
[Human]SARS-CoV-2   NKVVSTTTNI VTRCLNRV-- ---------- --------CT NYMPYFFTLL LQLC--TFTR STNSRIKASM PTT---IAKN TVKSVGKFCL
[Human]MERS-CoV     FKEFATRTFT ATTAVGSCIK SVVRH---LG VTKGILTGCF SFVKMLFMLP LAYF--SDSK LGTTEVKVSA LKTAGVVTGN VVKQCCTAAV
[Bat]BtCoV/HKU9     ------TTRV TTSLLGGLVT RSVRKTADFV RSTNPGSKCV GLLCLFYQLF MRFW--LLVK KP------PI VKVSGIIAYN TGCGVTTCVL
[Human]HCoV-229E    ---------- ---------- ---------- --------FV IFFTWLLSMF TLCK--TAVT TGDVKIMAKA PQRTGVVLKR SLKYNLKA--
[Avian]IBV          NKPNLERIFN IAKKAIVGSS VVTTQ----- --------CG KLIGKAATFI ADKVGGGVVR NITDSIKGLC GITRGHFERK MSPQFLKTLM
[Murine]MHV-A59     ---LLLRDEK QEFVAPKVVK AKAIA----- --------CY CAVKWFLLYC FSWI--KFNT DN------KV IYTTEVASKL TFKLCCLA--
[Bat]BtCoV/HKU3     GQTAVITSNC IKKCVQRV-- ---------- --------FS NYMPYVITLL FQLC--TFTK STNSRIKASL PTT---IAKN SVKSVAKLCL
[Bat]BtCoV/512/2005 ---------- ---------- ---------- --------II VLIVYLFSLL AICF--RALK KRDMKVMAGV PERTGIILKR SVKYNYKA--
[Human]HCoV-OC43    LNLREIKPAV NVVKAVRNKT SA-------- --------CF NFIKWLFVLL FGWI--KISA DN------KV IYTTEIASKL TCKLVALA--
[Human]HCoV-NL63    ---------- ---------- ---------- --------IV LFLTWLLSMF SLLR--TSIM KHDIKVIAKA PKRTGVILTR SFKYNIRSAL
```

```
                    1891
[Feline]FCoV        ---VRQLNKP ALWRYIKLVL ------LLIA LY---HFFYL FVSIPVVHK- ---------- ---------- -------LAC SGSVQAYSNS
[Porcine]TGEV       ---MRQLNKP SVWRYAKLVL ------LLIA IY---NFFYL FVSIPVVHK- ---------- ---------- -------LTC NGAVQAYKNS
[Human]HCoV-HKU1    --LKNAFQTF RWSIFIKGFL ------VVAT VF---LFWFN FLYINVIFSD FYLPNISVFP IFVGRIVMWI KATFGLVTIC DFYSKLGVGF
[Bat]BtCoV/HKU4     YMLLGKFKRV DWKATLRLFL -LL---CTTI LL-LSSIYHL VLFNQVLSSD VMLEDATGIL AIYKE----V RSYLGIRTLC DGLVVEYRNT
[Human]SARS-CoV     DAGINYVKSP KFSKLFTIAM WLL---LLSI CL-GSLICVT AAFGVLLSN- ---------- ---------- ---FGAPSYC NGVRELYLNS
[Human]SARS-CoV-2   EASFNYLKSP NFSKLINIII WFL---LLSV CL-GSLIYST AALGVLMSN- ---------- ---------- ---LGMPSYC TGYREGYLNS
[Human]MERS-CoV     DLSMDKLRRV DWKSTLRLLL -ML---CTTM VL-LSSVYHL YVFNQVLSSD VMFEDAQGLK KFYKE----V RAYLGISSAC DGLASAYRAN
[Bat]BtCoV/HKU9     NYLRSRCGNI SWSRLLKLLR YMLYIWFVWT CLTICGVWLS EPYAPSLVT- ---------- --------RF KYFLGIVMPC DYVLVNETGT
[Human]HCoV-229E    ---SAAVLKS KWWLLAKFTK LLL---LIYT LY--SVVLLC VRFGPF---- ---------- ---------- -------NFC SETVNGYAKS
[Avian]IBV          FFLF-YFLKA SVKSVVASYK TVLCKVVLAT LL---IVWFV YTSNPVMFT- ---------- ------GIRV LDFLFEGSLC GPYKDYGKDS
[Murine]MHV-A59     --FKNALQTF NWSVVSRGFF ------LVAT VF---LLWFN FLYANVILSD FYLPNIGPLP TFVGQIVAWF KTTFGVSTIC DFYQVTDLGY
[Bat]BtCoV/HKU3     DVCINYVKSP KFSKLFTIVM WLL---LLSI CL-GSLTYVT AVLGVCLSS- ---------- ---------- ---LGVPSYC DGVRELYINS
[Bat]BtCoV/512/2005 ---LKFFFRL KFQYIKVFLK FSL---VLYT LY--ALMFMF IRFTPVGTP- ---------- ---------- --------IC KRYTDGYANS
[Human]HCoV-OC43    --FKNAFLTF KWSMVARGAC ------IIAT IF---LLWFN FIYANVIFSD FYLPKIGFLP TFVGKIAQWI KNTFSLVTIC DLYSIQDVGF
[Human]HCoV-NL63    -----FVIKQ KWCVIVTLFK FLL---LLYA IY--ALVFMI VQFSPFNS-- ---------- ---------- -------LLC GDIVSGYEKS

                    1981
[Feline]FCoV        S-FVKSEVCG N-SILCKACL ASYDELADFD HLQVS----- -WDYKSDPLW NRVIQLSYFI FLAVFGNNYV RCLLMYFVSQ YLNLWLSYFG
[Porcine]TGEV       S-FIKSAVCG N-SILCKACL ASYDELADFQ HLQVT----- -WDFKSDPLW NRLVQLSYFA FLAVFGNNYV RCFLMYFVSQ YLNLWLSYFG
[Human]HCoV-HKU1    T----SHFCN G-SFICELCH SGFDMLDTYA AIDFVQYEVD R-RVLFDYVS LVKLIVELVI GYSLYTVWFY PLFCLIGLQL FTTWLPDLFM
[Bat]BtCoV/HKU4     S-FDVMEFCS NRSVLCQWCL IGQDSLTRYS ALQMLQTHIT SYVLNIDWIW ---FALEFFL AYVLYTSSFN VLLLVVTAQY FFAYTSAFVN
[Human]SARS-CoV     SNVTTMDFCE G-SFPCSICL SGLDSLDSYP ALETIQVTIS SYKLDLTILG ---LAAEWVL AYMLFTKFFY LLGLSAIMQV FFGYFASHFI
[Human]SARS-CoV-2   TNVTIATYCT G-SIPCSVCL SGLDSLDTYP SLETIQITIS SFKWDLTAFG ---LVAEWFL AYILFTRFFY VLGLAAIMQL FFSYFAVHFI
[Human]MERS-CoV     S-FDVPTFCA NRSAMCNWCL ISQDSITHYP ALKMVQTHLS HYVLNIDWLW ---FAFETGL AYMLYTSAFN WLLLAGTLHY FFAQTSIFVD
[Bat]BtCoV/HKU9     ---------- --GWLHHLCM AGMDSLD-YP ALRMQQHRYG S-PYNYTYIL ---MLLEAFF AYLLYTPALP IVGILAVLHL IVLYLPIPLG
[Human]HCoV-229E    N-FVKDDYCD G-SLGCKMCL FGYQELSQFS HLDVV----- -WKHITDPLF SNMQPFIVMV LLLIFGDNYL RCFLLYFVAQ MISTVGVFLG
[Avian]IBV          --FDVLRYCA D-DFICRVCL HDKDSLHLYK HAYSVEQVYK DAASGFIFNW NWLYLVFLIL FVKPVAGFVI ICYCVKYLVL NSTVLQTGVC
[Murine]MHV-A59     R----SSFCN G-SMVCELCF SGFDMLDNYD AINVVQHVVD R-RLSFDYIS LFKLVVELVI GYSLYTVCFY PLFVLIGMQL LTTWLPEFFM
[Bat]BtCoV/HKU3     SNVTTMDFCQ G-YFPCSVCL SGLDSLDSYP ALETIQVTIS SYKLDLTFLG ---LAAEWLL AYMLFTKFFY LLGLSAIMQA FFGYFASHFI
[Bat]BtCoV/512/2005 T-FDKNDYCG N--VLCKICL YGYEELSDFT HTRVI----- -WQHLKDPLI GNILPLFYLV FLIIFGGFFV RIGITYFIMQ YINAAGVALG
[Human]HCoV-OC43    K----NQYCN G-SIACQFCL AGFDMLDNYK AIDVVQYEAD R-RAFVDYTG VLKIVIELIV SYALYTAWFY PLFALISIQI LTTWLPELFM
[Human]HCoV-NL63    T-FNKDIYCG N-SMVCKMCL FSYQEFNDLD HTSLV----- -WKHIRDPIL ISLQPFVILV ILLIFGNMYL RFGLLYFVAQ FISTFGSFLG

                    2071
[Feline]FCoV        YVKYSWFLHV V-------NF ESISVEFVII VVVFKAVLAL KHIFLPCNNP SCKTCSKIAR QTRIPIQVVV NGSMKTVYVH ANGTGKLCKK
[Porcine]TGEV       YVEYSWFLHV V-------NF ESISAEFVIV VIVVKAVLAL KHIVFACSNP SCKTCSRTAR QTRIPIQVVV NGSMKTVYVH ANGTGKFCKK
[Human]HCoV-HKU1    LETMHWLIRF IVFVANMLPA FVLLRFYIVV TAMYKVVGFI RHIVYGCNKA GCLFCYKRNC SVRVKCSTIV GGVIRYYDIT ANGGTGFCVK
[Bat]BtCoV/HKU4     WRAYNYIVSG LFFLVTHIPL HGLVRVYNFL ACLWFLRKFY SHVINGCKDT ACLLCYKRNR LTRVEASTIV CGTKRTFYIA ANGGTSYCCK
[Human]SARS-CoV     SNS--WLMWF IISIVQMAPV SAMVRMYIFF ASFYYIWKSY VHIMDGCTSS TCMMCYKRNR ATRVECTTIV NGMKRSFYVY ANGGRGFCKT
[Human]SARS-CoV-2   SNS--WLMWL IINLVQMAPI SAMVRMYIFF ASFYYVWKSY VHVVDGCNSS TCMMCYKRNR ATRVECTTIV NGVRRSFYVY ANGGKGFCKL
[Human]MERS-CoV     WRSYNYAVSS AFWLFTHIPM AGLVRMYNLL ACLWLLRKFY QHVINGCKDT ACLLCYKRNR LTRVEASTVV CGGKRTFYIT ANGGISFCRR
[Bat]BtCoV/HKU9     NS---WLVVF LYYIIRLVPF TSMLRMYIVI AFLWLCYKGF LHVRYGCNNV ACLMCYKKNV AKRIECSTVV NGVKRMFYVN ANGGTHFCTK
[Human]HCoV-229E    YKETNWFLHF I-------PF DVICDELLVT VIVIKVISFV RHVLFGCENP DCIACSKSAR LKRFPVNTIV NGVQRSFYVN ANGGSKFCKK
[Avian]IBV          FLD--WFVQT VF------SH FNFMGAGFYF WLFYKIYIQV HHILY-CKDV TCEVCKRVAR SNRQEVSVVV GGRKQIVHVY TNSGYNFCKR
[Murine]MHV-A59     LETMHWSARL FVFVANMLPA FTLLRFYIVV TAMYKVYCLC RHVMYGCSKP GCLFCYKRNR SVRVKCSTVV GGSLRYYDVM ANGGTGFCTK
[Bat]BtCoV/HKU3     SNS--WLMWF IISIVQMAPV SAMVRMYIFF ASFYYVWKSY VHIMDGCTSS TCMMCYKRNR ATRVECTTIV NGVKRSFYVY ANGGRGFCKA
[Bat]BtCoV/512/2005 YQDNVWLLHL L-------PF NSMGNIIVVA FIVTRILLFL KHVLFGCDKP SCIACSKSAK LTRVPLQTIL QGVTKSFYVN ANGGKKFCKK
[Human]HCoV-OC43    LSTLHWSFRL LVALANMLPA HVFMRFYIII ASFIKLFSLF KHVAYGCSKS GCLFCYKRNR SLRVKCSTIV GGMIRYYDVM ANGGTGFCSK
[Human]HCoV-NL63    FHQKQWFLHF V-------PF DVLCNEFLAT FIVCKIVLFV RHIIVGCNNA DCVACSKSAR LKRVPLQTII NGMHKSFYVN ANGGTCFCNK
```

```
                    2161
[Feline]FCoV        HNFYCKNCDS YGFDHTFICD EIVRDLSNSI KQTVYATDRS YQEVTKVECT DGFYRFYV-- ----GEEFTA YDYDVKHKKY SSQEVLK---
[Porcine]TGEV       HNFYCKNCDS YGFENTFICD EIVRDLSNSV KQTVYATDRS HQEVTKVECS DGFYRFYV-- ----GDEFTS YDYDVKHKKY SSQEVLK---
[Human]HCoV-HKU1    HQWNCFNCHS FKPGNTFITV EAAIELSKEL KRPVNPTDAS HYVVTDIKQV GCMMRLFY-- ----DRDGQR VYDDVDASLF VDINNLL---
[Bat]BtCoV/HKU4     HNWNCVECDT AGVGNTFICT EVANDLTTTL RRLIKPTDQS HYYVDSVVVK DAVVELHY-- ----NRDGSS CYERYPLCYF TNLEKLKFKE
[Human]SARS-CoV     HNWNCLNCDT FCTGSTFISD EVARDLSLQF KRPINPTDQS SYIVDSVAVK NGALHLYF-- ----DKAGQK TYERHPLSHF VNLDNLR---
[Human]SARS-CoV-2   HNWNCVNCDT FCAGSTFISD EVARDLSLQF KRPINPTDQS SYIVDSVTVK NGSIHLYF-- ----DKAGQK TYERHSLSHF VNLDNLR---
[Human]MERS-CoV     HNWNCVDCDT AGVGNTFICE EVANDLTTAL RRPINATDRS HYYVDSVTVK ETVVQFNY-- ----RRDGQP FYERFPLCAF TNLDKLKFKE
[Bat]BtCoV/HKU9     HNWNCVSCDT YTVDSTFICR QVALDLSAQF KRPIIHTDEA YYEVTSVEVR NGYVYCYF-- ----ESDGQR SYERFPMDAF TNVSKLH---
[Human]HCoV-229E    HRFFCVDCDS YGYGSTFITP EVSRELGNIT KTNVQPTGPA YVMIDKVEFE NGFYRLYS-- ----CETFWR YNFDITESKY SCKEVFK---
[Avian]IBV          HNWYCRNCDD YGHQNTFMSP EVAGELSEKL KRHVKPTAYA YHVVDEACLV DDFVNLKYKA ATPGKDSASS AVKCFSVTDF LKKAVFL-KE
[Murine]MHV-A59     HQWNCLNCNS WKPGNTFITH EAAADLSKEL KRPVNPTDSA YYSVTEVKQV GCSMRLFY-- ----ERDGQR VYDDVNASLF VDMNGLL---
[Bat]BtCoV/HKU3     HNWNCLNCDT FCAGSTFISD EVARDLSLQF KRPINPTDQS AYVVDSVTVK NGALHLYF-- ----DKAGQK TYERHPLSHF VNLDNLR---
[Bat]BtCoV/512/2005 HNFFCVDCDS YGYGCTFIND VIAPELSNVT KLNVIPTGPA TIIIDKVEFS NGFYYLYS-- ----GSTFWK YNFDITEAKY ACKDVLK---
[Human]HCoV-OC43    HQWNCIDCDS YKPGNTFITV EAALDLSKEL KRPIQPTDVA YHTVTDVKQV GCSMRLFY-- ----DRDGQR IYDDVNASLF VDYSNLL---
[Human]HCoV-NL63    HNFFCVNCDS FGPGNTFING DIARELGNVV KTAVQPTAPA YVIIDKVDFV NGFYRLYS-- ----GDTFWR YDFDITESKY SCKEVLK---

                    2251
[Feline]FCoV        TMFLLD---- -DFIVYN-PS GSSLASVRNV CVYFSQLIGR PIKIVNSELL EDL--SVDFK GALFNAKKNV IKNSFNVDVS ECKNL-----
[Porcine]TGEV       SMLLLD---- -DFIVYS-PS GSALANVRNA CVYFSQLIGK PIKIVNSDLL EDL--SVDFK GALFNAKKNV IKNSFNVDVS ECKNL-----
[Human]HCoV-HKU1    HSKVKV-VPN LYVVVVE--S DADRANFLNA VVFYAQSLYR PILLVDKKLI TTACNGISVT QIMFDVYVDT FMSHFDVDRK SFNNFVNIAH
[Bat]BtCoV/HKU4     VCKTPTGIPE HNFLIYDTND RGQENLARSA CVYYSQVLCK PMLLVDVNLV TTVGDSREIA IKMLDSFINS FISLFSVSRD KLEKLINTAR
[Human]SARS-CoV     ANNTKGSLPI -NVIVFDGKS KCDESASKSA SVYYSQLMCQ PILLLDQALV SDVGDSTEVS VKMFDAYVDT FSATFSVPME KLKALVATAH
[Human]SARS-CoV-2   ANNTKGSLPI -NVIVFDGKS KCEESSAKSA SVYYSQLMCQ PILLLDQALV SDVGDSAEVA VKMFDAYVNT FSSTFNVPME KLKTLVATAE
[Human]MERS-CoV     VCKTTTGIPE YNFIIYDSSD RGQESLARSA CVYYSQVLCK SILLVDSSLI TSVGDSSEIA TKMFDSFVNS FVSLYNVTRD KLEKLISTAR
[Bat]BtCoV/HKU9     YSELKGAAPA FNVLVFDATN RIEENAVKTA AIYYAQLACK PILLVDKRMV GVVGDDATIA RAMFEAYAQN YLLKYSIAMD KVKHLYSTAL
[Human]HCoV-229E    NCNVLD---- -DFIVFN-NN GTNVTQVKNA SVYFSQLLCR PIKLVDSELL STL--SVDFN GVLHKAYIDV LRNSFGKDLN ANMSL-----
[Avian]IBV          ALKCEQ-ISN DGFIVCNTQS AHALEEAKNA AIYYAQYLCK PILILDQALY EQL-VVEPVS KSVIDKVCSI LSSIISVDTA ALNYK-----
[Murine]MHV-A59     HSKVKG-VPE THVVVVE--N EADKAGFLGA AVFYAQSLYR PMLMVEKKLI TTANTGLSVS RTMFDLYVDS LLNVLDVDRK SLTSFVNAAH
[Bat]BtCoV/HKU3     ANNTKGSLPI -NVIVFDGKS KCEESAAKSA SVYYSQLMCQ PILLLDQALV SDVGDSTEVS VKMFDAYVDT FSATFSVPME KLKALVATAH
[Bat]BtCoV/512/2005 NCNILT---- -DFVVFN-NS GSNVTQVKNA CVYFSQLLCK PIKLVDSALL ASL--NVDFS ANLHKAFVEV LSNSFGKDLS NCSNM-----
[Human]HCoV-OC43    HSKVKS-VPN MHVVVVE--N DADKANFLNA AVFYAQSLFR PILMVDKNLI TTANTGTSVT ETMFDVYVDT FLSMFDVDKK SLNALIATAH
[Human]HCoV-NL63    NCNVLE---- -NFIVYN-NS GSNITQIKNA CVYFSQLLCE PIKLVNSELL STL--SVDFN GVLHKAYVDV LCNSFFKELT ANMSM-----

                    2341
[Feline]FCoV        ---------- ------EECY KLCN----LD VTFSTFEMAI NNAHRFGILI TDRSFNNFWP SKIKPGSSGV SAMDIGKCMT FDAKIVNAKV
[Porcine]TGEV       ---------- ------DECY RACN----LN VSFSTFEMAV NNAHRFGILI TDRSFNNFWP SKVKPGSSGV SAMDIGKCMT SDAKIVNAKV
[Human]HCoV-HKU1    ASLREGVQLE KVLDTFVGCV RKCC-SIDSD VETRFITKSM ISAVAAGLEF TDENYNNLVP TYLKSDN--I VAADLGVLIQ NGAKHVQGNV
[Bat]BtCoV/HKU4     DCVRRGDDFQ NVLKTFTDAA RGHA-GVESD VETTMVVDAL QYAHKNDIQL TTECYNNYVP GYIKPDS--I NTLDLGCLID LKAASVNQTS
[Human]SARS-CoV     SELAKGVALD GVLSTFVSAA RQGV--VDTD VDTKDVIECL KLSHHSDLEV TGDSCNNFML TYNKVEN--M TPRDLGACID CNARHINAQV
[Human]SARS-CoV-2   AELAKNVSLD NVLSTFISAA RQGF--VDSD VETKDVVECL KLSHQSDIEV TGDSCNNYML TYNKVEN--M TPRDLGACID CSARHINAQV
[Human]MERS-CoV     DGVRRGDNFH SVLTTFIDAA RGPA-GVESD VETNEIVDSV QYAHKHDIQI TNESYNNYVP SYVKPDS--V STSDLGSLID CNAASVNQIV
[Bat]BtCoV/HKU9     QQISSGMTVE SVLKVFVGST RAEAKDLESD VDTNDLVSCI RLCHQEGWEW TTDSWNNLVP TYIKQDT--L STLEVGQFMT ANAKYVNANI
[Human]HCoV-229E    ---------- ------AECK RALG----LS ISDHEFTSAI SNAHRCDVLL SDLSFNNFVS SYAKPEEK-L SAYDLACCMR AGAKVVNANV
[Avian]IBV          ---------- ------AGTL RDAL---LSI TKDEEAVDMA IFCHNHDVDY TGDGFTNVIP SYGIDTGK-L TPRDRGFLIN ADASIANLRV
[Murine]MHV-A59     NSLKEGVQLE QVMDTFIGCA RRKC-AIDSD VETKSITKSV MSAVNAGVDF TDESCNNLVP TYVKSDT--I VAADLGVLIQ NNAKHVQANV
[Bat]BtCoV/HKU3     SELAKGVALD GVLSTFVSAA RQGV--VDTD VDTKDVIECL KLSHHSDIEV TGDSCNNFML TYNKVEN--M TPRDLGACID CNARHINAQV
[Bat]BtCoV/512/2005 ---------- ------NECR ESLG---LSD VPEEEFSAAV SEAHRYDVLI SDVSFNNLIV SYAKPEEK-L AVHDIANCMR VGAKVVNHNV
[Human]HCoV-OC43    SSIKQGTQIY KVLDTFLSCA RKSC-SIDSD VDTKCLADSV MSAVSAGLEL TDESCNNLVP TYLKSDN--I VAADLGVLIQ NSAKHVQGNV
[Human]HCoV-NL63    ---------- ------AECK ATLG----LT VSDDDFVSAV ANAHRYDVLL SDLSFNNFFI SYAKPEDK-L SVYDIACCMR AGSKVVNHNV
```

```
                      2431
[Feline]FCoV             LTQRGKSVVW LSQDFSTLSS TAQKVLVKTF VEEGVNFSLT FNAVGSDEDL PYERFTESVS ---AKSG-
[Porcine]TGEV            LTQRGKSVVW LSQDFAALSS TAQKVLVKTF VEEGVNFSLT FNAVGSDDDL PYERFTESVS ---PKSG-
[Human]HCoV-HKU1         AKVANISCIW FIDAFNQLTA DLQHKLKKAC VKTGLKLKLT FNKQEASVPI L----TTPFS ---LKGG-
[Bat]BtCoV/HKU4          MRNANGACVW NSGDYMKLSD SFKRQIRIAC RKCNIPFRLT TSKLRAADNI L----SVKFS ATKIVG--
[Human]SARS-CoV          AKSHNVSLIW NVKDYMSLSE QLRKQIRSAA KKNNIPFRLT CATTRQVVNV I----TTKIS ---LKGG-
[Human]SARS-CoV-2        AKSHNIALIW NVKDFMSLSE QLRKQIRSAA KKNNLPFKLT CATTRQVVNV V----TTKIA ---LKGG-
[Human]MERS-CoV          LRNSNGACIW NAAAYMKLSD ALKRQIRIAC RKCNLAFRLT TSKLRANDNI L----SVRFT ANKIVGG-
[Bat]BtCoV/HKU9          AKGAAVNLIW RYADFIKLSE SMRRQLKVAA RKTGLNLLVT TSSLKADVPC M----VTPFK ---IIG--
[Human]HCoV-229E         LTKDQTPIVW HAKDFNSLSA EGRKYIVKTS KAKGLTFLLT INENQAVTQI P----ATSIV ---AKQGA
[Avian]IBV               --KNAPPVVW KFSELIKLSD SCLKYLISAT VKSGVRFFIT KSGAKQVIAC H----TQKLL VE-KKAG-
[Murine]MHV-A59          AKAANVACIW SVDAFNQLSA DLQHRLRKAC SKTGLKIKLT YNKQEANVPI L----TTPFS ---LKGG-
[Bat]BtCoV/HKU3          AKSHNVSLVW NVKDYMSLSE QLRKQIRSAA KKNNIPFRLT CATTRQVVNV I----TTKIS ---LKGG-
[Bat]BtCoV/512/2005      LTKDNVPVVW LAKDFIALSE EARKYIVRTT KTKGINFMLT FNDRRMHLTI P----TISVA ---NKKG-
[Human]HCoV-OC43         AKIAGVSCIW SVDAFNQFSS DFQHKLKKAC CKTGLKLKLT YNKQMANVSV L----TTPFS ---LKGG-
[Human]HCoV-NL63         LIKESIPIVW GVKDFNTLSQ EGKKYLVKTT KAKGLTFLLT FNDNQAITQV P----ATSIV ---AKQGA
```
